# A Large Electroencephalogram Database of Freewill Reaching and Grasping Tasks for Brain Machine Interfaces

**DOI:** 10.1101/2025.05.09.653170

**Authors:** Bhoj Raj Thapa, John Boggess, Jihye Bae

## Abstract

Brain machine interfaces (BMIs) offer great potential to improve the quality of life for individuals with neurological disorders or severe motor impairments. Among various neural recording modalities, electroencephalogram (EEG) is particularly favorable for BMIs due to its noninvasive nature, portability, and high temporal resolution. Existing EEG datasets for BMIs are often limited to experimental settings that fail to address subjects’ freewill in decision making. We present a large EEG dataset, containing a total of 6808 trials, recorded from 23 healthy young adults (8 females and 15 males with an age range from 18 to 24 years) while performing reaching and grasping tasks, where the target object is freely chosen at their desired pace according to their own will. This EEG dataset provides a realistic representation of reaching and grasping movement, making it useful for developing practical BMIs.

## Background & Summary

Brain machine interfaces (BMIs) enable communication between the BMI user and external devices, such as prosthetic limbs or a computer cursor. They have provided potential benefits for people with motor impairments, including spinal cord injuries [1], stroke [2], and amyotrophic lateral sclerosis [3]. Electroencephalogram (EEG)-based BMI is one of the most commonly considered forms of BMI primarily due to its noninvasive nature and high portability [4–10]. In addition, as EEG allows high temporal resolution with millisecond precision to capture neural dynamics, it has been favored for real time BMI applications where timely responses to user’s commands are essential [11]. Recent studies in EEG-based BMIs have shown great potential in various applications demonstrating their effectiveness [12, 13], including cursor control, speller systems [14, 15], wheelchair control [16], and robotic arm control [17]. For example, over 90% accuracy was achieved in 1- and 2-dimensional cursor control using EEG-based BMIs [18, 19], and a high information transfer rate of 325 bits/min was reported in EEG-based speller systems [20]. Similarly, the wheelchair control was successfully implemented, enabling safe navigation along the straight and curved path with zero collisions, using a hybrid control strategy of motor imagery and steady-state visual evoked potentials [21]. Furthermore, robotic arm control in 4 directions has been implemented in four subjects, achieving an average accuracy of 85.45% [22].

However, traditional EEG-based BMIs fail to address subjects’ freewill to choose a target object in movement-related tasks. That is, conventionally, the initiation time of the movement execution and the determination of a target object are predetermined by the experimenter. Freewill movements are inherently different from passive movements, as freewill movements rely heavily on the supplementary motor area, whereas the passive movements primarily involve the dorsal premotor cortex [23]. In addition, the differences are also reflected in EEG-based neural correlates. For example, stronger event-related synchronization, particularly in the 15-25 Hz frequency band in C3-Cz bipolar EEG, has been reported when self-paced paradigm is allowed, compared to the cue-based paradigm in a right-thumb switch pressing task [24]. Similarly, freewill movements exhibited higher peaks of movement-related potentials in the Cz channel for both shoulder abduction (12*µ*V vs. 6.5*µ*V cued) and finger abduction (9.5*µ*V vs. 6.2*µ*V cued) [25]. EEG recordings that allow subjects to freely choose when to move and which target to select can provide unique neural patterns, different from typical cue-based tasks where the movement execution time and decision-making of the target object are made by the experimenter. To the best of our knowledge, there is only one publicly available EEG dataset addressing freewill movement [26]. This data contains EEG recordings from 2 subjects when they were freely pressing one of two keys on a keyboard.

The majority of EEG-based BMIs rely heavily on motor imagery without having actual physical execution. As Supplementary Table summarizes [26–36], two commonly considered motor imagery strategies are a) imagining different body parts, such as hands, feet, and tongue, [26–30] and b) imagining the actual movement execution [26, 31, 32, 34–36]. The Supplementary Table 1 provides a list of 20 different datasets categorized based on experimental paradigms reported in 14 studies. Among the currently available 13 motor imagery EEG datasets, 6 of them correspond to a) imagining different body parts, and the remaining 7 to b) imagining actual movement execution. Imagining different body parts has often been preferred in EEG-based BMIs because it produces distinct neural patterns, particularly in primary and supplementary motor cortex [37, 38], however, this approach results in a less intuitive interaction for the BMI user [39]. For example, the left hand, right hand, both feet, and tongue imagery can be matched to 4 directions (left, right, up, and down, respectively) to control a cursor [28]. Associating both feet imagery with upward movement and tongue imagery with downward movement is counterintuitive. Although imagining the actual movement execution can be more straightforward from the user’s perspective, distinguishing among different movement-related intentions from EEG can be challenging, requiring sophisticated neural representations combining movement complexity [40], goal [41], trajectory [42], and speed [43].

**Table 1:** Participant Demographics and Data Summary.

| Subject ID | Age [year] | Sex | fs [Hz] | Day Number | Sessions | Trials | Number of Trials |  |  |  |
| --- | --- | --- | --- | --- | --- | --- | --- | --- | --- | --- |
|  |  |  |  |  |  |  | Tgt1 | Tgt2 | Tgt3 | Tgt4 |
| AWL419 | 21 | F | 250 | 1 | 6 | 170 | 37 | 47 | 40 | 46 |
|  |  |  |  | 2 | 7 | 184 | 38 | 56 | 39 | 51 |
| BCX014 | 18 | F | 250 | 1 | 7 | 206 | 51 | 75 | 34 | 46 |
| BER853 | 19 | M | 250 | 1 | 6 | 156 | 31 | 40 | 42 | 43 |
|  |  |  |  | 2 | 6 | 191 | 42 | 59 | 44 | 46 |
| CNE421 | 21 | M | 250 | 1 | 4 | 117 | 27 | 40 | 22 | 28 |
|  |  |  |  | 2 | 7 | 194 | 38 | 57 | 48 | 51 |
| EIK361 | 22 | M | 250 | 1 | 4 | 118 | 26 | 33 | 28 | 31 |
|  |  |  |  | 2 | 4 | 118 | 28 | 32 | 28 | 30 |
|  |  |  |  | 3 | 4 | 117 | 27 | 32 | 26 | 32 |
| EPQ831 | 18 | M | 250 | 1 | 5 | 143 | 31 | 39 | 34 | 39 |
|  |  |  |  | 2 | 6 | 170 | 37 | 44 | 39 | 50 |
| GKE482 | 21 | F | 250 | 1 | 3 | 89 | 19 | 26 | 22 | 22 |
|  |  |  |  | 2 | 3 | 90 | 19 | 25 | 23 | 23 |
|  |  |  |  | 3 | 4 | 119 | 28 | 31 | 30 | 30 |
| GRI855 | 22 | M | 250 | 1 | 3 | 90 | 22 | 22 | 21 | 25 |
|  |  |  |  | 2 | 4 | 119 | 27 | 30 | 30 | 32 |
| HIQ863 | 19 | F | 250 | 1 | 4 | 111 | 22 | 35 | 25 | 29 |
|  |  |  |  | 2 | 6 | 175 | 37 | 52 | 37 | 49 |
| HKE021 | 23 | M | 250 | 1 | 3 | 76 | 19 | 20 | 17 | 20 |
|  |  |  |  | 2 | 4 | 112 | 25 | 31 | 30 | 26 |
| HLQ983 | 18 | F | 250 | 1 | 6 | 173 | 35 | 50 | 48 | 40 |
|  |  |  |  | 2 | 7 | 207 | 50 | 58 | 50 | 49 |
| JWL109 | 19 | M | 250 | 1 | 5 | 146 | 32 | 42 | 32 | 40 |
|  |  |  |  | 2 | 7 | 203 | 45 | 58 | 54 | 46 |
| LLE031 | 20 | F | 1000 | 1 | 4 | 117 | 27 | 31 | 29 | 30 |
|  |  |  |  | 2 | 4 | 110 | 28 | 27 | 29 | 26 |
|  |  |  |  | 3 | 3 | 110 | 26 | 28 | 27 | 29 |
| NVB749 | 23 | M | 250 | 1 | 6 | 175 | 41 | 48 | 41 | 45 |
|  |  |  |  | 2 | 7 | 200 | 45 | 58 | 45 | 52 |
| OQE073 | 19 | M | 1000 | 1 | 3 | 85 | 17 | 25 | 15 | 28 |
|  |  |  |  | 2 | 5 | 116 | 27 | 35 | 23 | 31 |
|  |  |  |  | 3 | 3 | 91 | 17 | 29 | 19 | 26 |
| PJE146 | 20 | M | 250 | 1 | 6 | 165 | 37 | 54 | 37 | 37 |
|  |  |  |  | 2 | 6 | 170 | 38 | 52 | 38 | 42 |
| PYW942 | 22 | M | 250 | 1 | 3 | 86 | 20 | 24 | 20 | 22 |
| QPD964 | 21 | M | 250 | 1 | 5 | 136 | 25 | 37 | 33 | 41 |
|  |  |  |  | 2 | 6 | 179 | 39 | 48 | 42 | 50 |
| SOR746 | 24 | M | 250 | 1 | 3 | 88 | 21 | 26 | 21 | 20 |
|  |  |  |  | 2 | 4 | 115 | 31 | 32 | 24 | 28 |
| TRI071 | 21 | M | 250 | 1 | 6 | 155 | 44 | 40 | 38 | 33 |
|  |  |  |  | 2 | 5 | 143 | 31 | 50 | 37 | 25 |
| VNW394 | 22 | F | 250 | 1 | 6 | 174 | 41 | 50 | 33 | 50 |
|  |  |  |  | 2 | 6 | 173 | 44 | 48 | 38 | 43 |
| WKQ963 | 21 | M | 250 | 1 | 4 | 113 | 26 | 29 | 26 | 32 |
|  |  |  |  | 2 | 3 | 89 | 22 | 22 | 21 | 24 |
|  |  |  |  | 3 | 3 | 88 | 22 | 23 | 21 | 22 |
| ZLO981 | 23 | F | 250 | 1 | 6 | 169 | 40 | 47 | 37 | 45 |
|  |  |  |  | 2 | 6 | 167 | 40 | 45 | 42 | 40 |
| Mean | 20.74 |  |  | 2.13 | 4.86 | 138.94 | 31.47 | 39.63 | 32.22 | 35.61 |
| Std | 1.72 |  |  | 0.54 | 1.39 | 38.70 | 9.00 | 12.66 | 9.41 | 10.00 |
| Total |  |  |  | 49 | 238 | 6808 | 1542 | 1942 | 1579 | 1745 |

We provide EEG datasets recorded during actual movement execution. Motor imagery of movement execution and actual movement execution exhibit similar neural dynamics and pathways [44]. A comparative study of alpha and beta band desynchronization during both conditions across 22 subjects revealed that desynchronization was consistently observed in both conditions, with 18 subjects during actual movement execution and 11 subjects during motor imagery [45]. Furthermore, when comparing movement execution and movement imagination for finger movements, no statistical difference was found in latency and amplitude of lateraized readiness potential, which is one of the components of the movement-related potentials [46]. The overlap between motor imagery and actual movement execution is further supported by evidence showing that both activate motor-related brain areas, including the premotor cortex, the dorsolateral prefrontal cortex, and the supplementary area [47]. Therefore, EEG data recorded from actual movement execution can be utilized to characterize motor imagery of actual movements, which can help develop BMIs. Among the 20 available EEG datasets, only 7 EEG datasets [26, 31, 32, 34, 48–50] are recorded while performing an actual upper limb movement, and only 1 dataset [26] is recorded while performing freewill movement execution (Supplementary Table 1).

Large, standardized, and openly accessible datasets promote the development of reliable signal processing, analysis methods, and decoder models for EEG-based BMIs [51–53]. Currently, there exists only one EEG dataset available for a freewill upper limb task [26]. This dataset includes only 2 subjects performing a freewill key-press task. In comparison, our dataset includes 23 subjects while performing a freewill reaching and grasping task that mirrors common real-world movements. Furthermore, our dataset includes a total of 6808 trials. This substantial increase in data volume further bolsters the importance of our dataset for studying voluntary motor control in tasks that mimic common activities in daily life.

In addition, our dataset includes a simultaneous EEG and electrooculogram (EOG) recording that enables the investigation of efficient artifact removal strategies. Only 9 publicly available EEG datasets include EOG [26, 30, 31, 35, 48]. Eye movement artifacts can contaminate scalp EEG, particularly in the frontal and prefrontal regions [39, 54]. The available EOG allows for the removal of ocular artifacts, such as blinks and saccades, that interfere with EEG signal quality [55–58]. Effective removal of artifacts can improve the overall signal-to-noise ratio in EEG signals [59, 60]. Using the cleaned EEG signal after EOG correction (based on the simultaneously recorded EOG), the classification accuracy for sleep stage classification provides an improvement of 8.03% [61]. Similarly, the EOG correction method using simultaneously recorded EOG also helped in the classification of motor imagery tasks, where it improved the classification accuracy by an average of 2.5% across 9 subjects [62].

Furthermore, we provide continuous EEG data, containing both movement planning [63, 64] and execution phases [44, 65]. This dataset allows capturing transitions between movement phases (before and after the movement onset) and non-movement phases (resting or idle) where a larger temporal context is useful to develop BMIs [66–68]. This is also helpful to develop practical BMIs, as continuous EEG input enables the simulation of online BMI implementation, allowing the system to dynamically interpret user intentions as they arise [69]. Additionally, continuous data allows us to inspect how neural responses adapt and change during motor tasks with time, which is often overlooked in segmented datasets, losing broader temporal context. Among 20 available EEG datasets as shown in Supplementary Table 1, 16 provide continuous data, including multiple consecutive trials, offering flexibility for studying real-time BMI applications.

In this paper, we publish a large freewill EEG dataset [70]. Unique features of our dataset are listed below:

- **Freewill Movements**: During the experiment, subjects were allowed to decide the movement initiation time and target object by their own will to complete a reaching and grasping task. This specific experimental design provides the opportunity to investigate unique neural dynamics associated with freewill.
- **Actual Movement Execution**: During the experiment, subjects moved their dominant arm and hand to complete a reaching and grasping task. Understanding the specific neural dynamics related to actual movement can help design effective BMIs.
- **Large Dataset**: This dataset includes a total of 49 EEG recordings from 23 healthy subjects with a total of 6808 trials. This far exceeds the publicly available freewill finger movement dataset, which contains EEG recordings from 2 subjects [26].
- **Available EOG**: Synchronously recorded 4 channel EOG is provided. The available EOG signals can facilitate the investigation of ocular artifact removal strategies. Successful removal of ocular artifacts can improve the EEG quality, which can ultimately help to obtain a reliable neural decoder in BMI.
- **Raw Continuous Recording**: The provided raw continuous data allows the implementation of various signal processing techniques and understanding of continuous neural dynamics associated with pre- and post-movement intentions.

## Methods

### Participants

23 healthy right-handed young adults participated in this study. The participants included 8 female and 15 male subjects aged between 18 and 24 years (20.74 ± 1.72 years). A total of 6808 trials were collected in 49 recording days. Table 1 summarizes the participant demographics, including age and sex, and collected EEG recording details, including the total number of trials per recording. All recordings were conducted in the Neural Interfaces and Signal Processing (NISP) lab housed in the Department of Electrical and Computer Engineering at the University of Kentucky. This study was conducted following the approved Institutional Review Board (IRB) protocol #57031. Participants went through a screening process to ensure the absence of neurological disorders, any ongoing medication that would potentially affects neural activities, and potential contradictions to the designed experimental paradigm. Before data collection, all participants were informed of the study’s purpose and procedures, and they provided written consent for data collection. At the start of each recording day, a detailed questionnaire was used to review the participant’s general health condition to ensure their eligibility to participate in the recording, including any change in medication and fatigue levels. To uphold confidentiality, all personal information was anonymized, and each participant was assigned a unique subject ID composed of a random combination of 3 letters followed by 3 digits (e.g., *AWL419*).

### Experimental Paradigm

During the experiment, the participant was comfortably seated on an office chair in front of a sturdy brown desk, wearing an EEG cap. Four different cups were placed on the table in front of the subject, and each cup was numbered in a counterclockwise manner from the participant’s perspective. Cups were named Tgt1, Tgt2, Tgt3, and Tgt4 (Figure 1) based on the order. The detailed measurements of the table and cup setup using transparent shelves positioned to hold the cups is shown in Figure 2. The cups Tgt1 and Tgt3 were filled with water, while Tgt2 and Tgt4 were empty. A designated red square and 2 white rectangles were marked on the table to guide participant’s movement. The red square indicates the final destination where the selected cup must be placed, and the white rectangles show where subjects are instructed to keep both hands while resting. To minimize visual disturbances, participants were seated facing a plain white cement wall.

**Figure 1.**
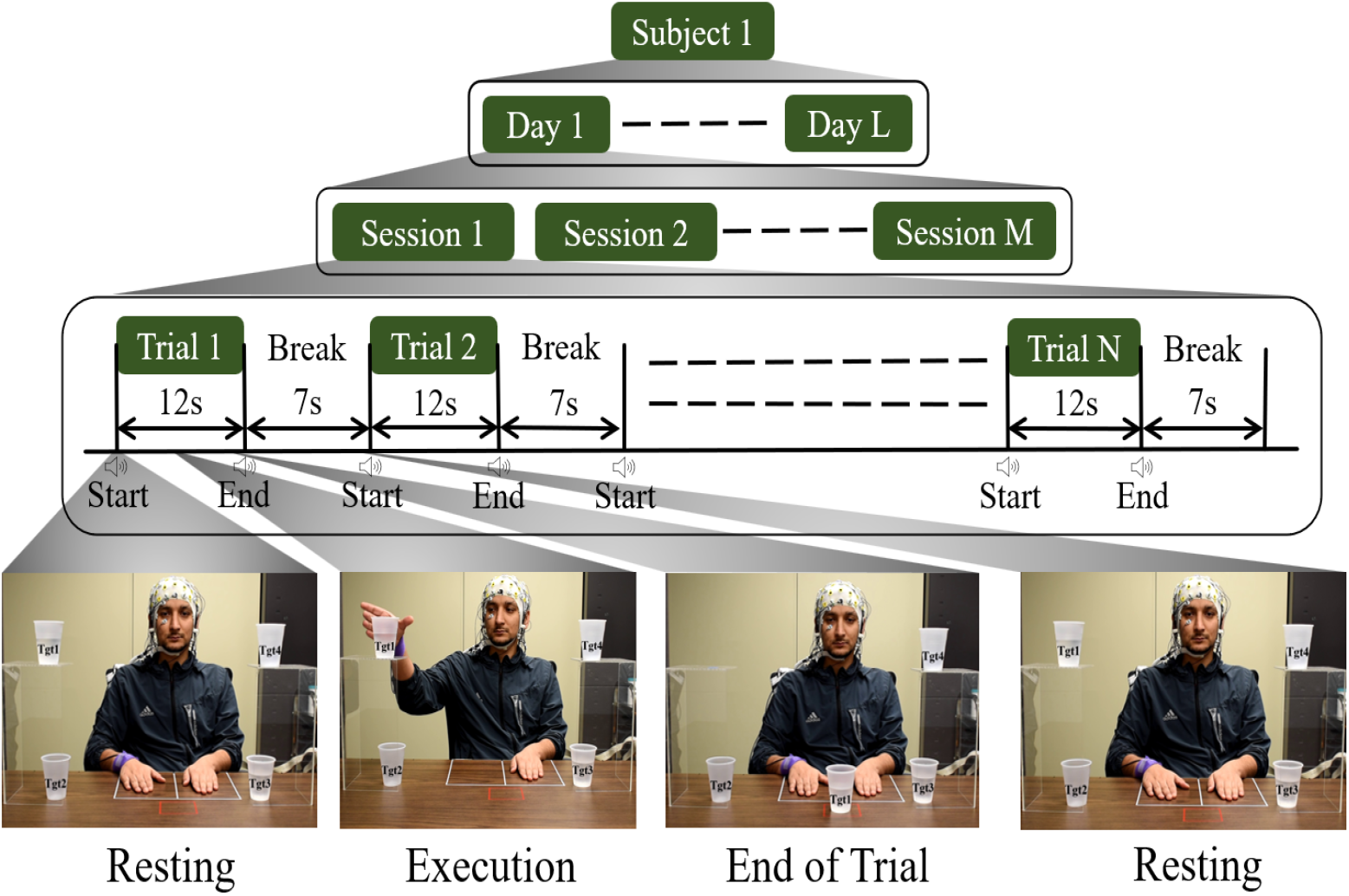
Experimental Paradigm of the Freewill Reaching and Grasping Task. In the experiment, each subject completes multiple recordings from different days; each recording consists of several sessions, and each session contains multiple trials. The top green block represents one subject as *Subject 1*, and the green blocks in the 2nd row indicate multiple recordings from different days, *Day 1* to *Day L*, where *L* represents the total number of recordings. The blocks at the 3rd row illustrate that each recording includes several sessions from *Session 1* to *Session M*), and the 4th row shows that each session consists of multiple trials, from *Trial 1* to *Trial N*. Each trial lasts 12 seconds, and it is followed by a 7-second break. Each trial begins with an audio cue, ‘start,’ in which the subject performs the freewill reaching and grasping task at any time within the 12 seconds using only their dominant hand (right). In each trial, the subject performs reaching and grasping one of the 4 cups (Tgt1, Tgt2, Tgt3 or Tgt4) with their own will. After a trial ends, a 7-second break starts with an ‘end’ audio cue, during which the subject returns the selected cup to its original place. The bottom row panel provides visual snapshots of the subject’s actions across different stages: *Resting* where the subject maintains a relaxed position before the starting a trial, *Execution* where the subject performs the freewill reaching and grasping task any time within the 12-second period, *End of Trial* where subject completes the task, and *Resting* where the subject relaxes after placing the selected cup in its original pace. The pictures include the image of the first author, and he consented to include the identifiable picture, per his preference.

**Figure 2.**
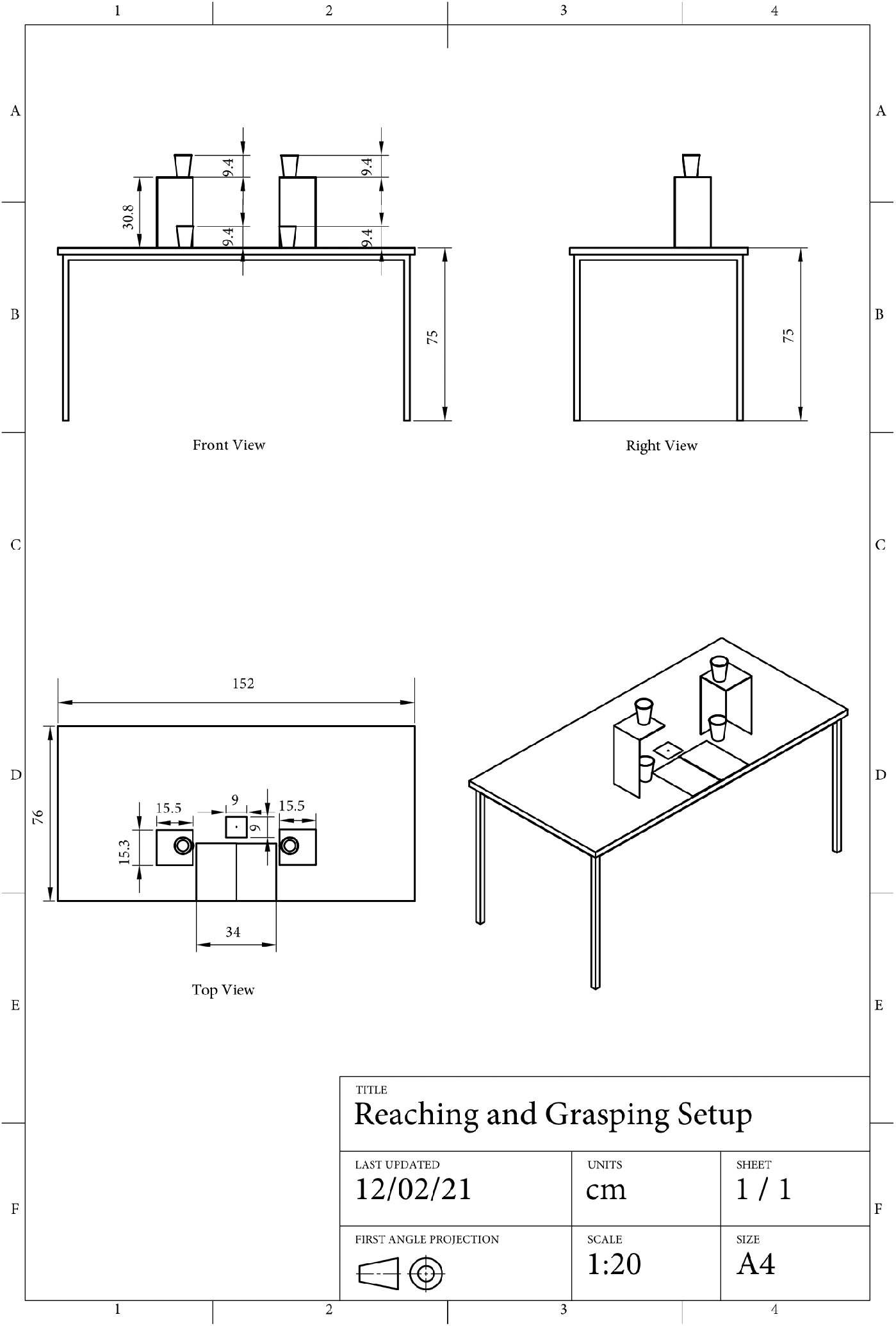
Orthographic Projection of the Experimental Setup. This figure represents an orthographic projection of the experimental setup, illustrating its front (top left), right side (top right), and top (bottom left) views. All dimensions are in centimeters, and the drawing follows first angle projection conventions with a scale of 1:20.

Participants typically completed 2 or 3 recordings on different days except for 2 participants (*BCX014* and *PYW942*), who completed only one recording, due to their preference not to continue with further recordings. Each recording includes multiple sessions, ranging from 3 to 7, where each session contains a block of trials, which typically includes 30 trials, ranging from 20 to 35 trials. Between sessions, a break of around 2 to 5 minutes was provided mainly to allow participants to rest. During the break, the experimenters also ensured that the subjects felt comfortable continuing the recording and addressed any questions brought by the subjects. The number of sessions per recording and the number of trials per session varied on the basis of several factors, including the time available after the preparation of electrodes for the recording and the participants’ fatigue and willingness to continue.

An illustration of the described experimental paradigm is provided in Figure 1, and an example video *SOR746_ExampleVideo*.*mov* is provided in the following directory *data/experiment_video/*. Each trial starts with a ‘start’ audio cue and ends with an ‘end’ audio cue, and lasts a total of 12 seconds. Both ‘start’ and ‘end’ cues are played through 2 speakers (Bose Companion 2 Series III Multimedia Speaker System for PC), which are placed on the left side of the subjects, with the volume maintained at approximately 65 dB. Participants were instructed to complete a reaching and grasping task at their own pace and will, within the 12-second trial duration. The subject is instructed to wait for around 1 to 2 seconds after hearing the start cue to avoid event-related potentials (ERPs) in EEG caused by the auditory stimulus. During a trial, the subject freely selects one of the four cups. If they chose a cup with water (Tgt1 or Tgt3), they were instructed to take a sip of water and then place the cup in the designated red square on the table. When the subject selects an empty cup (Tgt2 or Tgt4), the subject is instructed to place the cup directly onto the red square. They were instructed to perform the task as naturally as possible using only their dominant hand (right), while maintaining a uniform speed and consistent hand and arm postures throughout the entire recording. A 7-second break starts with the ‘end’ cue and ends with the next ‘start’ cue for the next trial, and the sequence continues until the session ends. During the 7-second break, the subject is instructed to place the selected cup back to its original location and then put their hand back to the resting area within the white rectangle on the table. During the resting phase, the participants were asked to relax their muscles to minimize the artifacts caused by voluntary muscle contraction and were encouraged to maintain the same initial posture and position during the entire recording sessions.

### Data Acquisition System

During the experiment, 31 channel EEG, 4 channel EOG, 1 audio, and 3 accelerometer signals were simultaneously recorded. The EEG signal was recorded using a gel-based active electrode system (actiCAP snap cap with actiCHamp Plus amplifier, Brain Products). This includes 31 EEG electrodes (Fp1, Fp2, F7, F3, Fz, F4, F8, FT9, FC5, FC1, FC2, FC6, FT10, T7, C3, C4, T8, TP9, CP5, CP1, CP2, CP6, TP10, P7, P3, Pz, P4, P8, O1, Oz, and O2) with Cz as the reference and GND as the ground (Figure 3 (a)). 4 additional gel-based active electrodes were placed around the eyes to record eye movements (Figure 3 (b)). 2 EOG electrodes were placed beside the left (channel name of EOGL) and right eye (channel name of EOGR) to capture horizontal eye movements, and the remaining 2 EOG electrodes were placed under (channel name of EOGU) and above the right eye (channel name of EOGR) to record vertical eye movements. Each recording was conducted by 2 trained experimenters, and the electrode placement was always reviewed by an expert.

**Figure 3.**
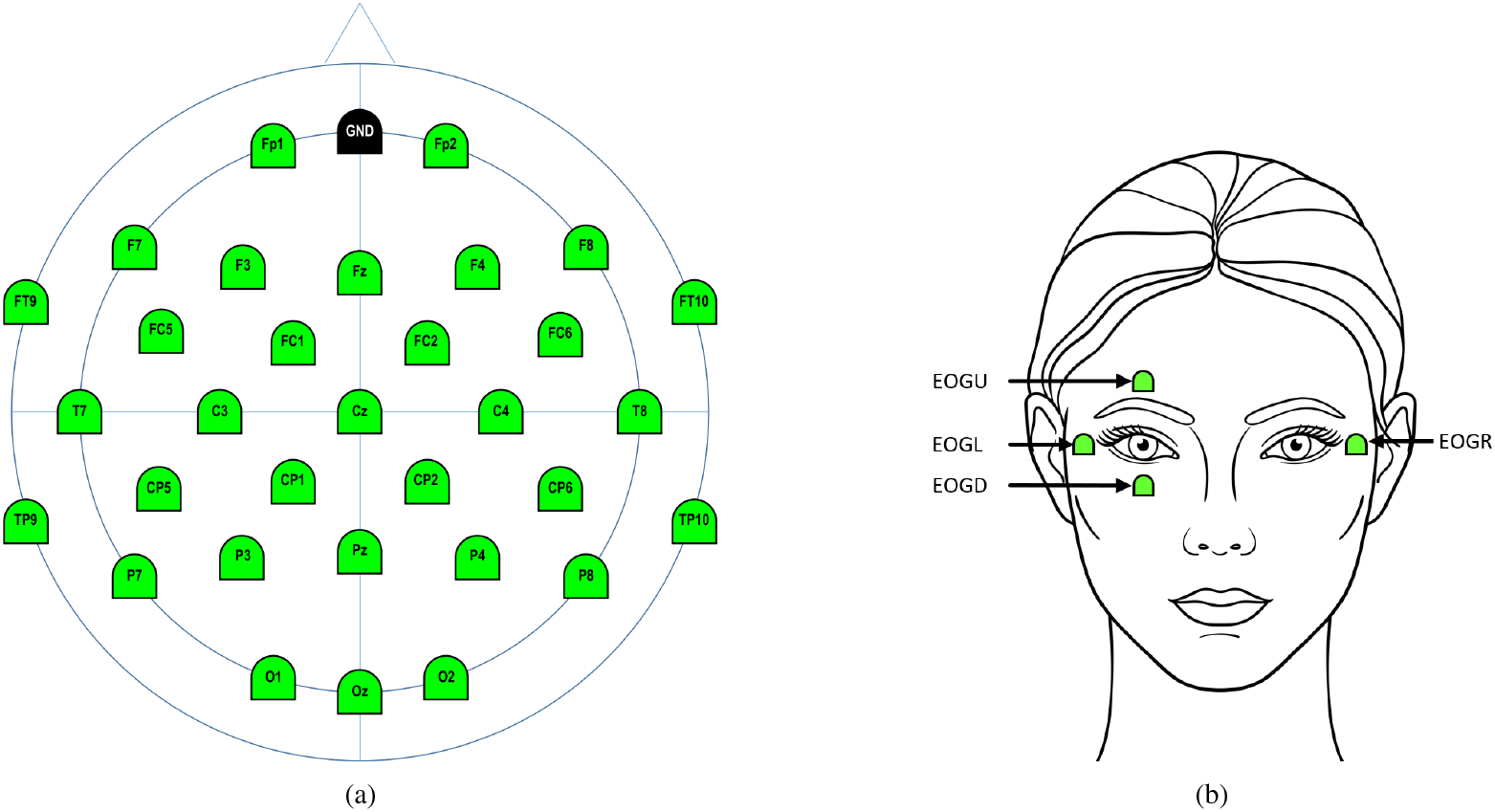
EEG and EOG Electrode Locations. (a) EEG and (b) EOG configurations. In (a), green colored electrodes display all electrodes used for the recording, and the black colored electrode shows where the ground is located. In (b), green colored electrodes indicate where each EOG electrode was placed for the recording. EOGL (left) and EOGR (right) electrodes are placed to capture horizontal eye movements, while EOGU (upper) and EOGD (lower) electrodes are for capturing vertical eye movements.

A custom program script (MATLAB Version 2022a, MathWorks) was used to play the timed ‘start’ and ‘end’ audio cues, and the audio signal (channel name of TRIG) was connected to the amplifier’s auxiliary port (StimTrak, Brain Products). A lightweight 3D accelerometer sensor (Acceleration Sensor Amplifier Module S/N ACC20050249, Brain Products) was placed left to the distal end of the ulna to record the tri-axial acceleration (channel names of X, Y, and Z) from the hand movements, and each channel was connected to the amplifier’s auxiliary port. The accelerometer values were used primarily to detect the movement onset.

As all signals are connected to the same amplifier, the 31 channel EEG, 4 channel EOG, 1 audio signal, and 3 accelerometer signals were synchronized and sampled at the same sampling frequency (Figure 4). A sampling frequency of 1000 Hz was used for 2 of the subjects (*LLE031* and *OQE073*), while the signals of the remaining 21 subjects were sampled at 250 Hz. The amplifier was connected to a workstation (Dell PC with 64 GB RAM, NVIDIA Quadro P2000 GPU, and Intel Xeon Silver 4114 CPU), running Windows 11, and all signals were collected using commercial recording software (BrainVision Recorder Version 1.23.0003, Brain Products).

**Figure 4.**
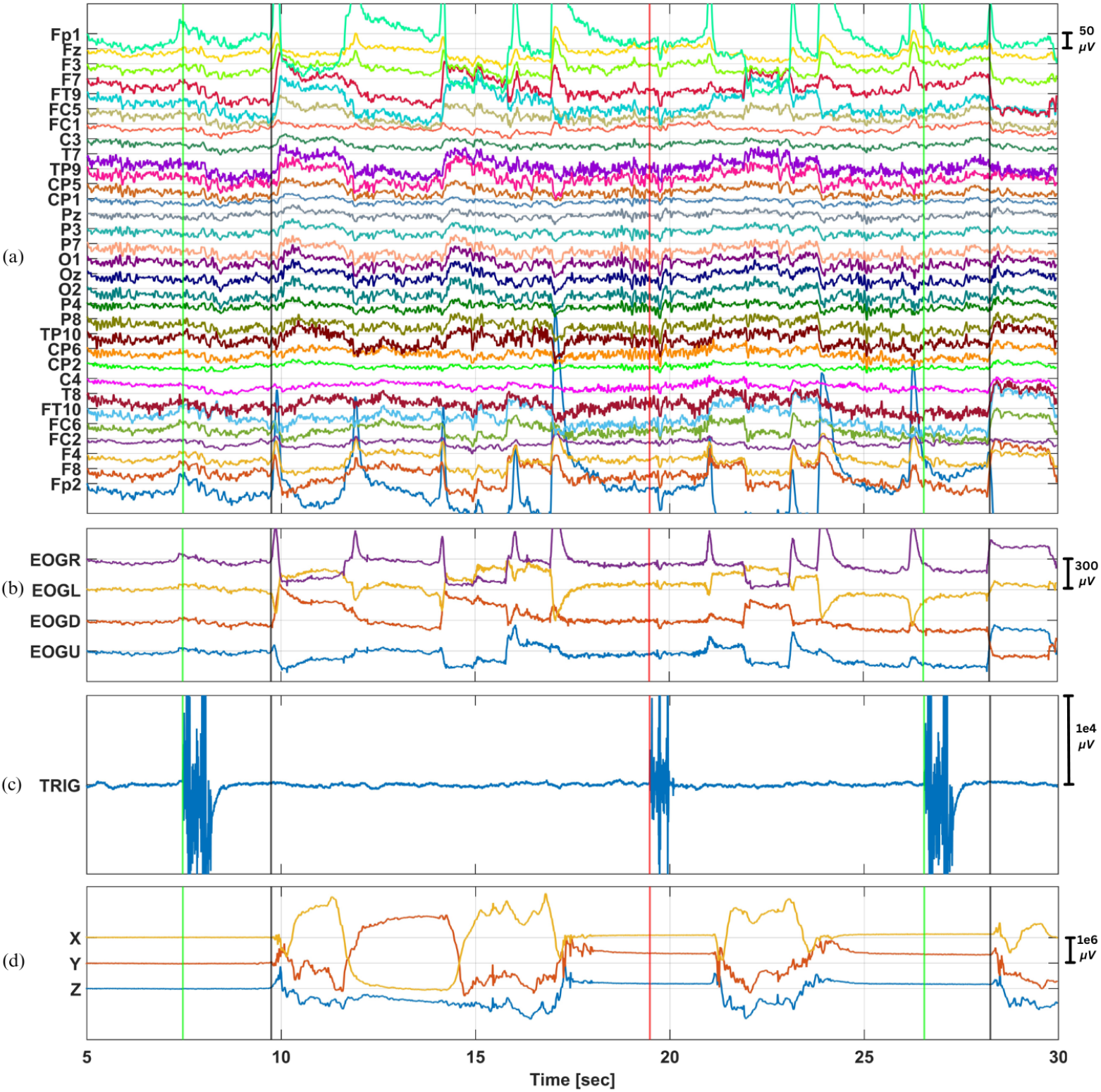
An Example of Recorded Data. (a) 31 channel EEG after applying 0.1-30 Hz band-pass filter, (b) 4 channel raw EOG, (c) 1 channel audio (TRIG), and (d) 3 channel raw accelerometer (X, Y, and Z) data. The x-axis displays the time in seconds. The y-axis labels denote the channel names. The green vertical lines mark the ‘start’ of each trial, while the red vertical line indicates the ‘end’ of the trial, which was determined based on the TRIG channel. The black vertical line represents the movement onset, where the subject started making the movement, detected based on X, Y, and Z channels. All signals (EEG, EOG, TRIG, X, Y, and Z) are time synchronized. The data from *AWL419_Day1*.*mat* was extracted from session 2, visualizing from 5 to 30 seconds of recordings.

## Data Records

### Distribution for use

The dataset for the freewill reaching and grasping task is provided as a .*zip* file, named *Freewill_EEG_Reaching_Grasping*.*zip*. It can be accessed from the Figshare data repository service (https://doi.org/10.6084/m9.figshare.28632599) [70]. The dataset includes raw EEG data files, an example experiment video, and supporting code to generate audio cues played during the experiment and to reproduce all reported results.

### Data Directory Structure

The data directory structure depicted in Figure 5 explains how the dataset is organized. There are 2 main directories,*data* and *src* and a *README*.*md* file. The 2 main directories contain several subdirectories. Details about these directories are provided in the following subsections. The *README*.*md* file includes explanations on how to navigate the data directories, process the raw data, and any necessary details for running the provided scripts in the *src* directory.

**Figure 5.**
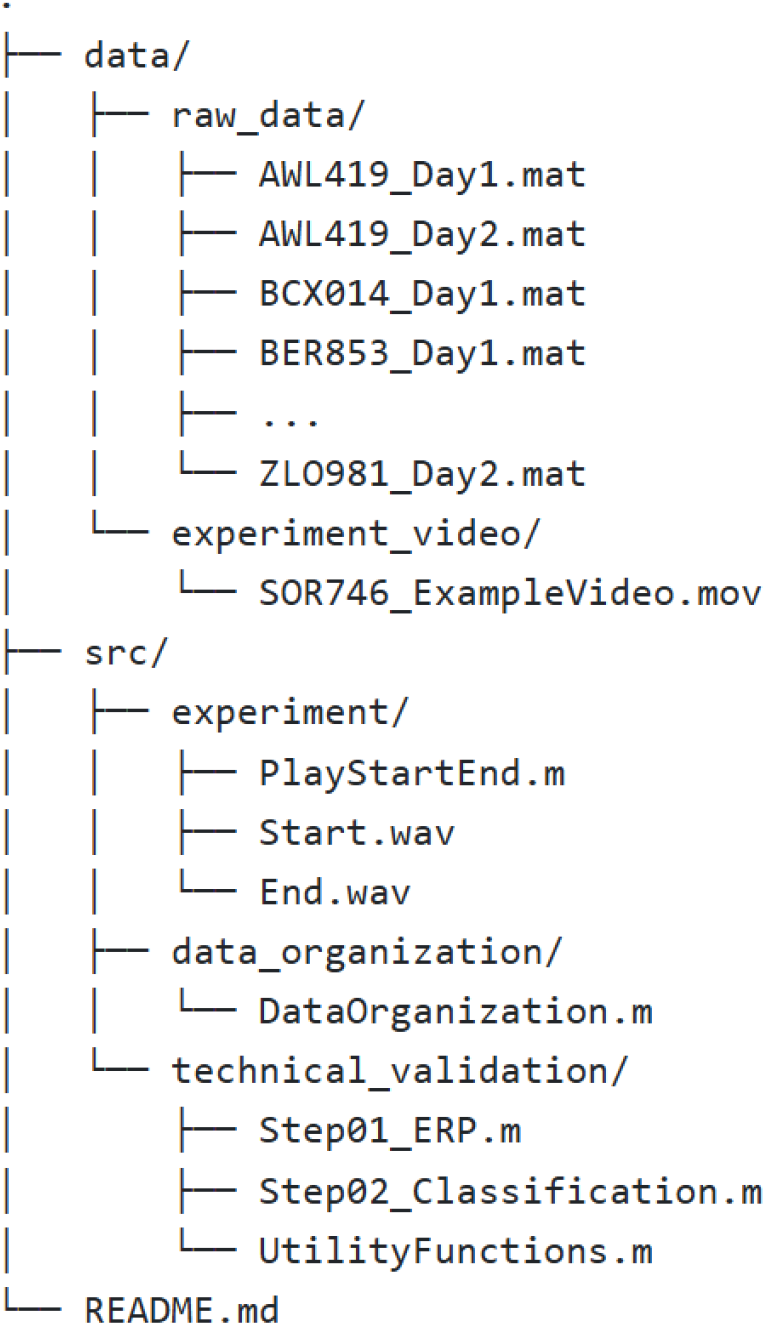
Directory Structure of Shared Files. The shared *Freewill_EEG_Reaching_Grasping*.*zip* file follows a specific structure, containing 3 main branches of *data/, src/*, and *README*.*md*. The *data/* directory contains all recorded data, with *data/raw_data* storing all 49 recordings and *data/experiment_video* providing 1 example video. The *src/* directory includes all MATLAB scripts to conduct the experiment and to generate reported results. *src/experiment* contains the scripts used during the experiment, *src/data_organization* provides the scripts to organize the data with detected the trial start and end and movement onset time points, and *src/technical_validation* contains scripts to generates all reported results, described in section Technical Validation. Finally, the *README*.*md* file includes a detailed description of the .*zip* file and its directory structure, helping efficient navigation and use of the dataset.

#### Main Directory 1: data

The *data* directory contains 2 subdirectories of *raw_data* and *experiment_video*.

- **Raw Data Directory, *data/raw_data/*** This directory contains 49 recordings from all 23 subjects in *SubjectID_DayL*.*mat* format. Each data file represents 1 recording from 1 subject. The file name follows the convention *SubjectID_DayL*.*mat*, where *SubjectID* represents subject’s unique identifier, and *DayL* indicates the day of recording. For example, for the first day of recording by a subject named *AWL419*, the file name *AWL419_Day1*.*mat* was assigned. Each file includes 39 channel raw data (31 channel EEG, 4 channel EOG (channel names EOGU, EOGD, EOGL, and EOGR), 1 audio signal (channel name TRIG), and 3 accelerometer signals (channel names X, Y, and Z)), along with necessary details of the data, such as sampling frequency and channel labels. Details on the data file are provided in the following section EEG Data Structure.
- **Example Video Directory, *data/experiment_video/*** This directory contains 1 example video file, *SOR746_ExampleVideo*.*mov*. This video is recorded while a subject *SOR746* is conducting 1 session of the experiment. This video displays the overall experimental paradigm as described in Experimental Paradigm section. The subject *SOR746* consented to include the identifiable video per the subject’s preference.

#### Main Directory 2: src

This *src* directory contains MATLAB code to conduct the experiment, process the data, and generate the reported results.

- **Experiment Code, *src/experiment/*** This subdirectory includes 1 MATLAB script and 2 audio files used during the experiment. *PlayStartEnd*.*m* is a MATLAB script used to play the audio cues to guide the start and end of each trial with the specific time intervals, and *Start*.*wav* and *End*.*wav* files that are called to play the audio cues to indicate the ‘start’ and ‘end’ of the trial, respectively.
- **Data Organization Code, *src/data_organization/*** This subdirectory contains 1 MATLAB script, *DataOrganization*.*m*, designed to structure a data table that contains raw data and key information of each trial. This script also processes the TRIG and XYZ channels to identify time points for the trial start and end and movement onset. The original recordings generated by the recording software (BrainVision Recorder Version 1.23.0003, Brain Products) are in .*eeg* format. These .*eeg* files were first exported as .*dat* files using a commercial software (BrainVision ANalyzer Version 2.2, Brain Products). In MATLAB, the raw data from these .*dat* files were then organized, and key information was saved in each *SubjectID_DayL*.*mat*. The *DataOrganization*.*m* detects the trial start and end time points in each trial based on the TRIG channel. In addition, after determining the trial start and end time points, this script detects the movement onset time point. Detailed explanations of these detection processes are provided in section Data Processing. Once these time points are identified, the script stores all necessary information in a *dataTable* cell array. Details on this data table are provided in EEG Data Structure section.
- **Technical Validation Code, *src/technical_validation/*** This subdirectory consists of 3 MATLAB scripts of *Step01_ERP*.*m, Step02_Classification*.*m*, and *UtilityFunctions*.*m*. They are designed to preprocess EEG signals and perform classification tasks, where *UtilityFunctions*.*m* are used for both *Step01_ERP*.*m* and *Step02_Classification*.*m* scripts. *Step01_ERP*.*m* processes the raw EEG to compute average event-related potentials (ERPs), visualizes ERPs, and stores the processed EEG as a new MATLAB structure field in the *SubjectID_DayL*.*mat* file to be used in the following classification step in (*Step02_Classification*.*m*). A detailed explanation of ERP computation and visualization processes is provided in section Event-Related Potential (ERP). The *Step02_Classification*.*m* script loads the processed EEG, constructs input features, applies a linear support vector machine (LSVM), and computes the performance accuracy. The *UtilityFunctions*.*m* script contains 4 functions: *getSubjectNames(), countDaysForSubject(), extract_epochs()*, and *plot_multichannelEEG()*. The *getSubjectNames()* function identifies unique subject IDs, and *countDaysForSubject()* counts the number of recordings available for each subject. These functions are particularly useful because the *data/raw_data/* directory contains 49 files, each including the subject ID and day number. By automating the process, these functions efficiently extract subject IDs and determine the number of recording days for each subject based on all the file names listed within the *data/raw_data/* directory. The *extract_epochs()* function creates time-locked EEG epochs from continuous signals based on the event marker, such as the movement onset. In addition, *plot_multichannelEEG()* function plots multichannel EEG data, such as the one shown in Figure 4 (a).

### EEG Data Structure

Each *SubjectID_DayL*.*mat* file in the *data/raw_data* directory provides a structured EEG dataset, containing 8 fields storing key information, as shown in Figure 6. This data file is compatible with both MATLAB and Python environments. The scripts available in section *Main Directory 2: src/technical_validation/* show how to load this *SubjectID_DayL*.*mat* file in MATLAB using the built-in *load()* function. For Python, one way to load this data file is to use *scipy*.*io*.*loadmat()* function from *SciPy* library [71]. Each data file contains 1 EEG recording from 1 subject. Detailed descriptions of each field are provided below:

**Figure 6.**
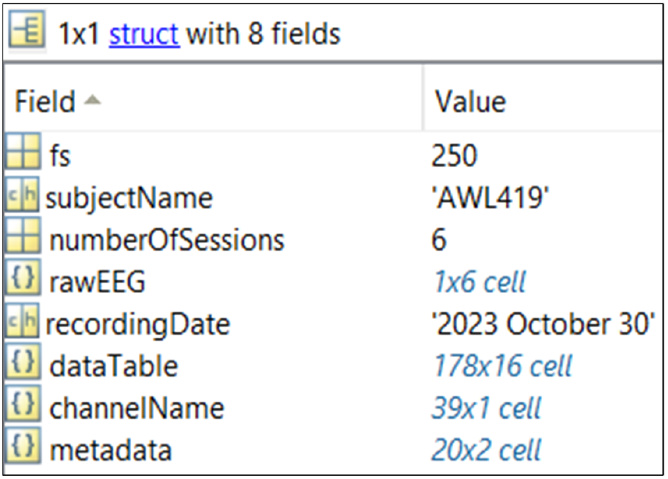
Structure of EEG Data File. Each *SubjectID_DayL*.*mat* file, contains 8 fields of sampling frequency (*fs*), subject’s ID (*subjectName*), total number of sessions in the recording (*numberOfSessions*), raw recorded data (*rawEEG*), the date of recording (*recordingDate*), key information of each trail including trial start and end time and movement onset time indices (*dataTable*), channel names of the recording (*channelName*), and the description of these fields (*metadata*). This example is from the data file *Data_AWL419_Day1*.*mat*. The same data structure is followed by all shared data files.

- ***fs***: sampling frequency of the recording in Hz.
- ***subjectName***: subject’s ID in a combination of 3 letters and 3 numerical numbers.
- ***numberOfSessions***: total number of sessions included within the data.
- ***rawEEG***: a [1 × *numberOfSessions*] cell array containing raw recordings. Each cell includes a [39 × total number of samples] matrix, containing raw EEG, EOG, TRIG, X, Y, and Z signals for a specific session. The 39 rows represent 31 EEG channels, 4 EOG channels, 1 TRIG channel, and 3 acceleration channels of X, Y, and Z. The cell array’s column number corresponds to the session number. That is, data recorded during the first session, Session 1, is stored in cell (1,1), for the 2nd session in cell (1,2), and so on.
- ***recordingDate***: recording date in long date format (e.g., 2023 October 30) on which the recording was conducted.
- ***dataTable***: a [(total number of trials+2) × 16] cell array, storing trial specific information for each trial. An example of *dataTable* is provided in Figure 7. The first row of the *dataTable* specifies the 16 column headers, the last row provides a sum over 5 key columns of *Tgt1, Tgt2, Tgt4, Tgt4* and *TrialDiscardIndex*, and the rows in between the first and the last rows provide trial specific information for each trial. Details on each column are as follows:
  - ***SessionNumber***: session number of the recording, where each session is composed of multiple trials.
  - ***TrialNumber***: the trial number within a corresponding session, indicated in *SessionNumber*.
  - ***Tgt1, Tgt2, Tgt3, Tgt4***: binary indicators showing the target cup reached during each trial by the subject. One of the 4 columns must have a value of ‘1’ representing the selected trial, and the rest must be ‘0.’ The target number is assigned in each cup as shown in Figure 1.
  - ***TgtID***: an integer value from 1 to 4 indicating the selected target number. Based on the binary values in *Tgt1* to *Tgt4*, 1 target number is assigned.
  - ***TrialDiscardIndex***: a binary flag for bad trials. ‘1’ indicates a bad trial to be discarded, and ‘0’ indicates to keep. Bad trials include the trials where the subject failed to complete the instructed movement or when external noise affects the subject’s task performance or signal quality.
  - ***TrialStartIndex***: positive integer values indicating at which sample index each trial starts. The index was detected using the method explained in the Trial Start and End Time section.
  - ***TrialEndIndex***: positive integer values indicating at which sample index each trial ends. The index was detected using the method explained in the Trial Start and End Time section.
  - ***AccStartIndex***: positive integer values indicating at which sample index movement onset is detected. Details on the detection approaches are explained in the Movement Onset Detection section.
  - ***AccStartIndexFlag***: a binary flag that indicates whether the movement onset detection was completed manually ‘1’ or automatically ‘0.’
  - ***TRIGSTARTDetectionFlag***: a binary indicator for whether trial start was detected manually ‘1’ or automatically ‘0.’
  - ***TRIGENDDetectionFlag***: a binary indicator specifying whether the trial end was detected manually ‘1’ or automatically ‘0.’
  - ***EEGFileNumber***: a unique 4-digit identifier linking the trial to its original .*eeg* file. It tracks the session order for each subject, starting at 0001 for the first session and incrementing by 1 for subsequent sessions (e.g., 0002, 0003, and so on). The numbering may not always be sequential due to discarded sessions.
  - ***Comments***: experimenter’s notes providing additional context for trials. This provides notes of any unusual or irregular events that happened during the experiment, such as unexpected subject’s movement, external interferences, or anomalies in the recorded signal. For example, ‘subject coughed’ is commented when subject coughed during the experiment, ‘accelerometer tangled in between the experiment’ for when the accelerometer connecting wire got tangled hindering the task performance, ‘long breathing’ for instances of heavy irregular breathing by the subject and ‘HF oscillations’ for noticeable high frequency oscillations in the data.
- ***channelName***: a [39 × 1] cell array, containing all channel names. The length of *channelName* matches the number of rows in each cell in *rawEEG*, as each corresponding row’s channel names for EEG, EOG, the audio, and accelerometer signals are listed.
- ***metadata***: A [20 × 2] cell array containing descriptions of the *SubjectID_DayL*.*mat* file, so that the data can be understood independently, making the *SubjectID_DayL*.*mat* a standalone file. The first row serves as a header, with ‘Name’ in the first column and ‘Description’ in the second column. Of the remaining 19 rows, 7 rows list the field names of the *SubjectID_DayL*.*mat* file, while the other 12 rows describe the 12 columns (*SessionNumber, TrialNumber, TgtID, dataTableDiscardIndex, TrialStartIndex, TrialEndIndex, AccStartIndex, AccStartIndexFlag, TRIGSTARTDetectionFlag, TRIGENDDetectionFlag, EEGFileNumber* and *Comments*) of the *dataTable* within the *SubjectID_DayL*.*mat* file.

**Figure 7.**
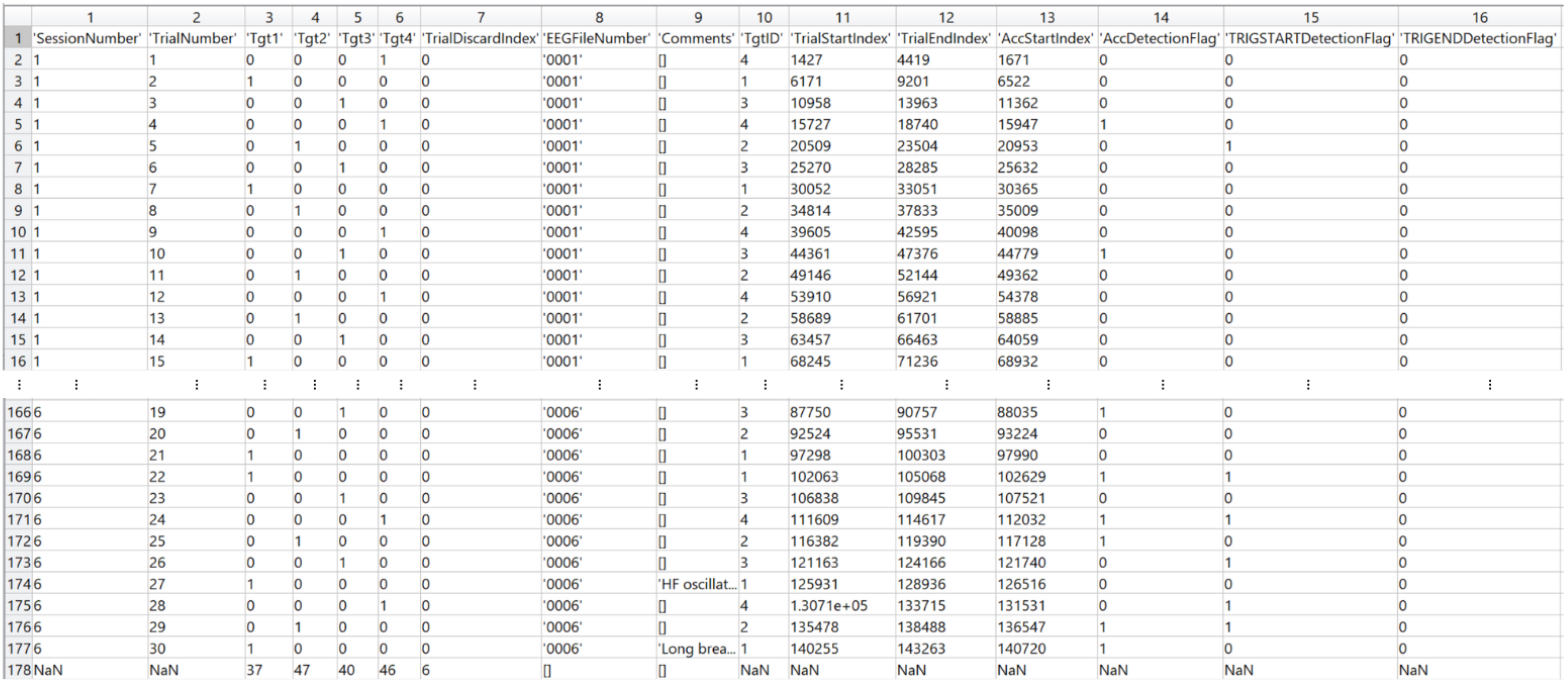
An Example of *dataTable*. This is an example of *dataTable* cell array which is a part of *SubjectID_DayL*.*mat* file. There are 16 columns, and each row consists of each trial’s details. The details of this cell array are explained in *dataTable*. This example is from the data file *Data_AWL419_Day1*.*mat*. The same data structure is followed by all shared data files.

### Data Processing

Detecting the trial start, trial end, and movement onset time in each trial is a critical step in processing the acquired data. Here, we provide a detailed process to detect these time points. The start and end times of each trial were identified based on the TRIG channel, and the movement onset time was determined by the X, Y, and Z channels.

### Trial Start and End Time

Figure 4 (c) displays an example of TRIG signal that is generated from the audio cues, where the vertical green and red lines indicate the start and end time of each trial. The trial start and end time points were determined by detecting the time points when the ‘start’ and ‘end’ cues began, respectively. These time points were identified using the *DataOrganization*.*m* script, located in *src/data_organization/* (see section Data Organization Code). A zero-phase high pass filter with a 1 Hz cutoff frequency was applied to the TRIG channel to remove the DC component. After a first order absolute difference was computed on the filtered TRIG signal, the first peak above a threshold was identified. The threshold was automatically calculated as twice the maximum amplitude within the first second of TRIG data from the recording, which contains mainly background noise. All automatically marked trial start and end time points were reviewed visually by a trained experimenter. In cases where the automatically detected start and end time points did not align with the beginning of each peak, the experimenter visually evaluated the pattern of cues in the TRIG signal and manually determined the correct start of the peak. Once one trial’s start and end time points were determined, the automatic detection algorithm sought the subsequent trial’s start and end time points after 7 seconds from the trial end time point to avoid false detections. This 7-second gap was used to ensure the minimum interval between cues, based on the experimental paradigm. The time indexes of the ‘start’ and ‘end’ cues are stored under *TrialStartIndex* and *TrialEndIndex* columns in *dataTable*, respectively.

### Movement Onset Time

Figure 4 (d) shows an example of the 3-dimensional acceleration signals (X, Y, and Z) used to detect movement onset, where the vertical black lines indicate the determined movement onset. The movement onset time for each trial was determined using the script named *DataOrganization*.*m* stored in *src/data_organization/* (see section Data Organization Code). The acceleration signals captured the movement onset within a trial, the time between ‘start’ and ‘end’ cues. First, a damaged channel that did not show significant change within a trial was discarded. We used a threshold of the standard deviation of the first 0.1 seconds to determine the damaged channel. After the valid axes were identified, the norm of all available axes was computed. Then, the norm was passed through a zero-phase low pass filter with a 10 Hz cutoff frequency. After the first order difference was computed on the smoothed norm signal, the threshold of 2000 was used to detect the movement onset. This threshold was determined by analyzing the histogram of the first order difference values from 4 randomly selected subjects. Movement onset was marked at the first sample where the difference exceeded the threshold within the time window between the ‘start’ and ‘end’ of the trial. All automatically detected movement onset time points were reviewed by the experimenters. For any trial where automatic detection was inaccurate, the time point was manually adjusted. Once finalized, the time index of the movement onset is stored under *AccStartIndex* in *dataTable*.

## Technical Validation

### Ocular Artifacts Correction

The applicability of using the 4 EOG channels to remove ocular artifacts is evaluated. Eye movement artifacts such as blinking and saccades can distort EEG. Simultaneously recorded 4 EOG channels (EOGU, EOGD, EOGL, and EOGR) along with the multichannel scalp EEG enable effective removal of ocular artifacts [72]. For example, EEG signals can be decomposed by independent component analysis (ICA), and independent components showing high similarity to EOG channels can be removed. After removing the selected independent components, the EEG signal can be reconstructed. ICA algorithms, such as FastICA [73], Extended InfoMax [74], and JADE [75], have been popularly used for ocular artifact removal due to their effectiveness [76]. In particular, Extended InfoMax has shown improved performance in removing biological artifacts, including eye movements in EEG, compared to FastICA and JADE. Extended Infomax achieved significantly lower residual artifact errors, particularly in the delta band for 5%, compared to 15% for FastICA and 12% for JADE, indicating minimal distortion of neural signals [77]. Extended InfoMax uses a flexible optimization framework to maximize mutual information, enabling a better separation of independent sources in complex data with mixed sub-Gaussian and super-Gaussian distributions [74, 78].

By using Extended Infomax, independent components are identified in the 31 channel raw EEG signals. The identified independent components were then compared with vertical and horizontal eye movements using the EOGD and EOGU channel pair for vertical activity, and the EOGL and EOGR channel pair for horizontal activity. Their similarity was measured using the sum of squared correlation (SSC), which quantifies how closely each independent component matches the EOG signals. A threshold of 15% of the total SSC was applied. That is, independent components contributing at least 15% of the total SSC score were flagged as representing ocular activity. The threshold of 15% was chosen because it consistently identified 2 key IC components corresponding to vertical and horizontal eye activities. After removing these two components, the EEG signal was reconstructed [79].

Figure 8 illustrates the effective use of the recorded EOG channels to remove the ocular artifacts on EEG signals using Extended Infomax. Before applying the ocular artifact correction, large fluctuations corresponding to vertical and horizontal eye movements are clearly visible in EEG channels, around 10 and 12 seconds (red circles in Figure 8 (a)). The highlighted EEG patterns are very similar to the dynamics of the recorded EOG channels, as especially, EOGR and EOGD channels show prominent eye movement activity (Figure 8 (b)). After the ocular artifact correction, these artifacts are substantially attenuated (Figure 8 (c)). In particular, significant changes in Fp1 and Fp2 channels are observed as the amplitudes of those channels were substantially reduced.

**Figure 8.**
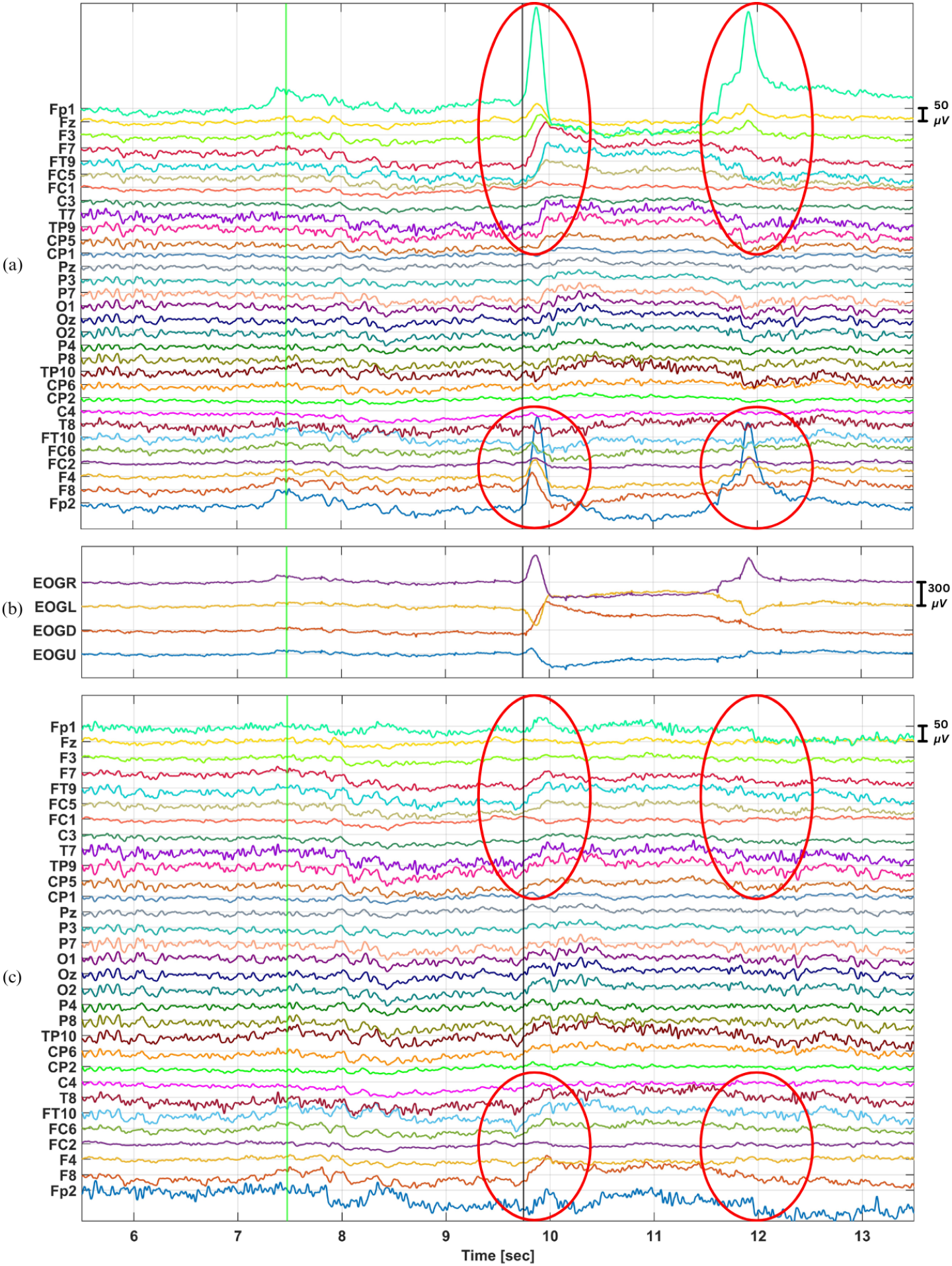
An Example of Ocular Artifact Correction. (a) 31 channel EEG after applying 0.1-30 Hz band-pass filter, (b) synchronized 4 channel EOG (raw), and (c) processed EEG after ocular artifact correction. The x-axis and y-axis are the time in seconds and channel names, respectively. The green and black vertical lines mark the start of the trial and movement onset, respectively. In (a), the red circled regions around 10-12 seconds highlight the predominant effects of ocular artifacts in EEG. In (b), the red circled areas emphasize the effective ocular artifact correction. This figure illustrates an example of EEG data of *AWL419_Day1*.*mat* extracted from session 2 from 5.5 to 13.5 seconds.

### Event-Related Potential (ERP)

To validate our dataset, we visualize average ERPs. Starting from movement onset, we display average ERPs with respect to the 4 different targets for 3 seconds in Figures 9, 10, and 11. First, EEG data from the 2 subjects (*LLE031* and *OQE073*) with a sampling rate of 1000 Hz was downsampled to 250 Hz to be consistent with the other 21 subjects. During downsampling, an anti-aliasing lowpass infinite impulse response (IIR) filter with a 112.5 Hz cutoff was used, followed by a cubic spline interpolation. Then, the EEG data with 250 Hz sampling frequency was bandpass filtered between 0.1 and 30 Hz using a zero-phase 4th order Butterworth IIR filter using the *filtfilt()* function in MATLAB. For each EEG channel, the bandpass filtered EEG of trials corresponding to the same target was averaged, and the standard deviation was computed to assess variability. Note that for this ERP computation, only good trials (*TrialDiscardIndex* = ‘0’) were included.

**Figure 9.**
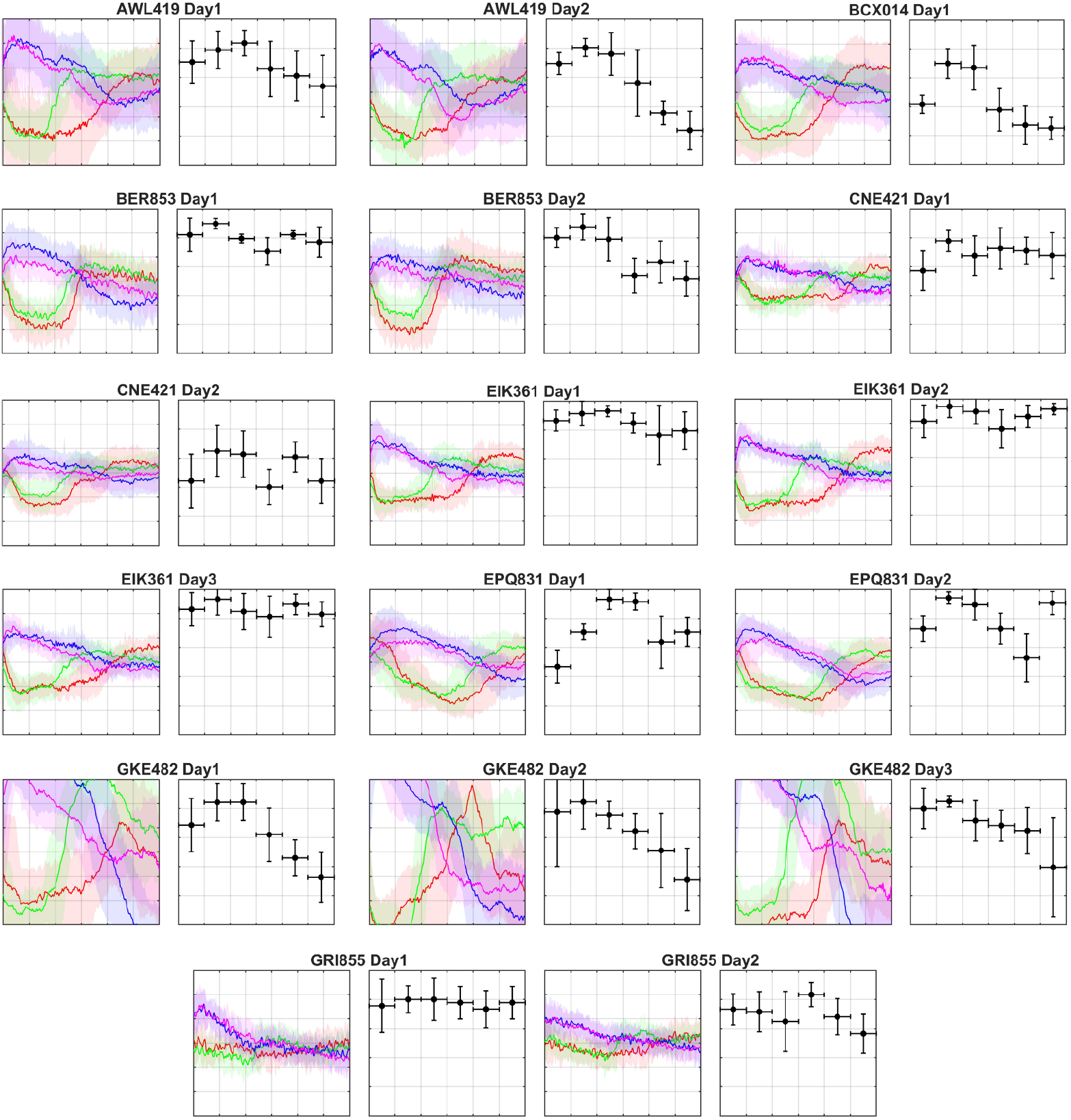
Average Event-Related Potentials (ERPs) and Neural Decoding Performance. Average ERPs and neural decoding results from EEG recordings, obtained from subjects AWL419 to GRI855, are displayed. Each subtitle represents the corresponding data file name. In each data set, average ERPs from the C3 channel are depicted on the left panel, and average neural decoding accuracies are displayed on the right panel. Each subfigure has x-axis ranges from 0 to 3 seconds relative to the movement onset (*t* = 0), with 0.5 grid interval. On the left panel, the average ERPs over all trials corresponding to the same target are displayed along with their standard deviation (shaded areas). The 4 targets of Tgt1, Tgt2, Tgt3, and Tgt4 are color coded in red, green, blue, and magenta, respectively. The y-axis shows an amplitude range of 30 ± *μ*V. The right panels represent average neural decoding accuracy over the 4 targets, when 0.5-second time window was used. The x-axis segment of each horizontal line indicates the time window used for the EEG extraction for LSVM implementation, whereas the y-axis value shows the average success rates. The y-axis ranges from 0.5 to 1 of accuracy, and each vertical line represents the standard deviation over 5 folds.

**Figure 10.**
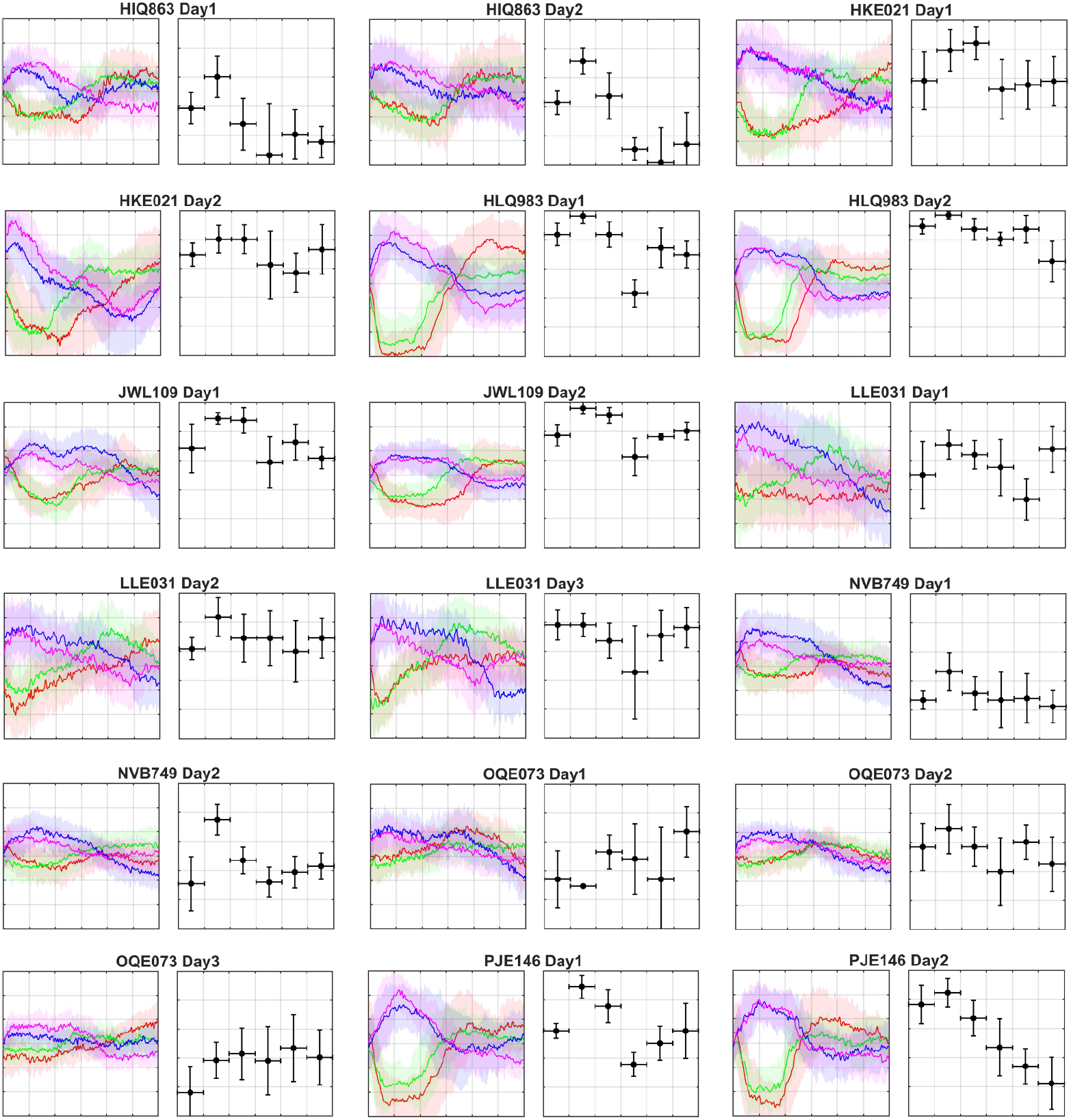
Average Event-Related Potentials (ERPs) and Neural Decoding Performance. Average ERPs and neural decoding results from EEG recordings, obtained from subjects HIQ863 to PJE146, are displayed. Each subtitle represents the corresponding data file name. In each data set, average ERPs from the C3 channel are depicted on the left panel, and average neural decoding accuracies are displayed on the right panel. Each subfigure has x-axis ranges from 0 to 3 seconds relative to the movement onset (*t* = 0), with 0.5 grid interval. On the left panel, the average ERPs over all trials corresponding to the same target are displayed along with their standard deviation (shaded areas). The 4 targets of Tgt1, Tgt2, Tgt3, and Tgt4 are color coded in red, green, blue, and magenta, respectively. The y-axis shows an amplitude range of 30 ± *μ*V. The right panels represent average neural decoding accuracy over the 4 targets, when 0.5-second time window was used. The x-axis segment of each horizontal line indicates the time window used for the EEG extraction for LSVM implementation, whereas the y-axis value shows the average success rates. The y-axis ranges from 0.5 to 1 of accuracy, and each vertical line represents the standard deviation over 5 folds.

**Figure 11.**
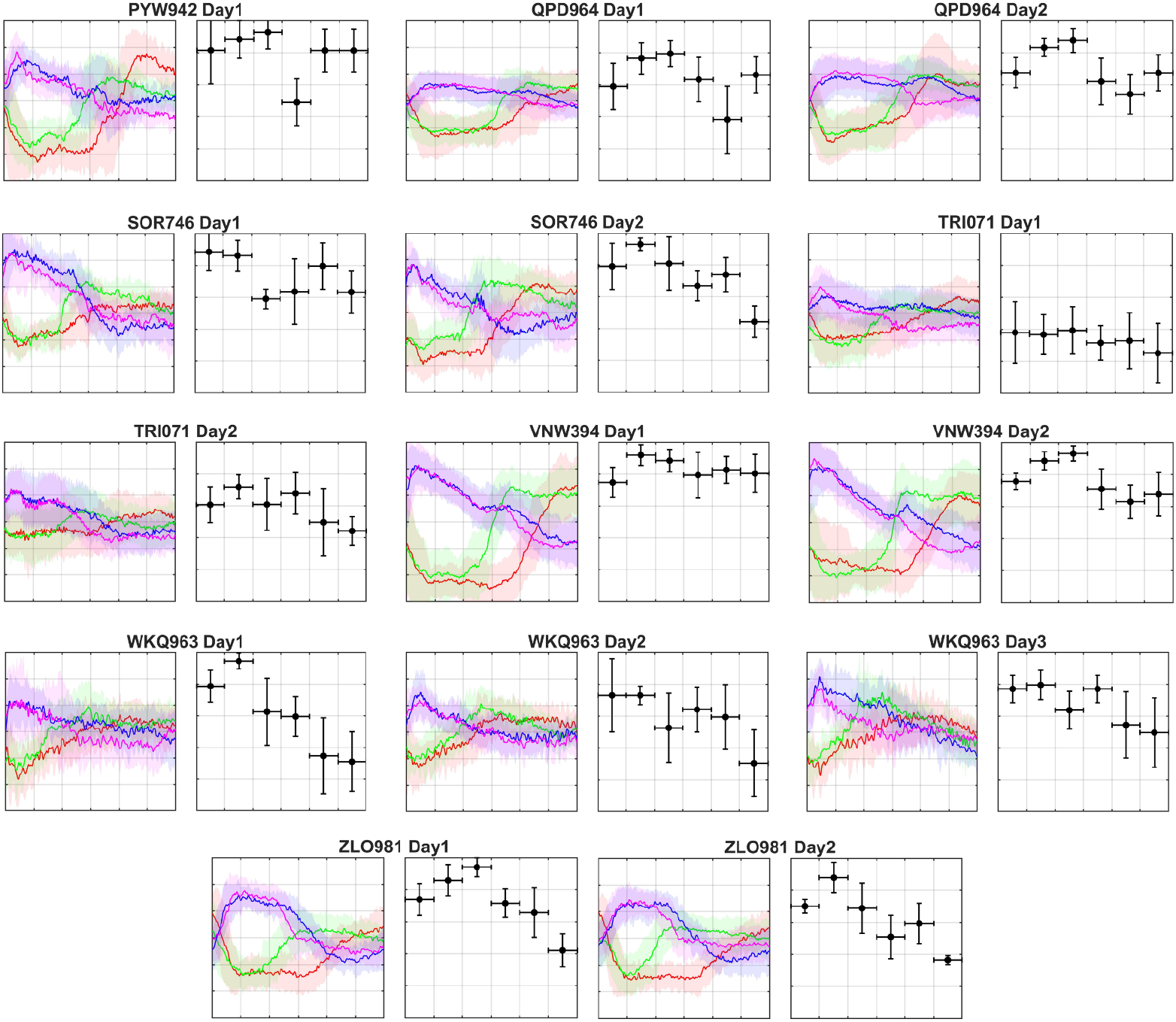
Average Event-Related Potentials (ERPs) and Neural Decoding Performance. Average ERPs and neural decoding results from EEG recordings, obtained from subjects PYW942 to ZLO981, are displayed. Each subtitle represents the corresponding data file name. In each data set, average ERPs from the C3 channel are depicted on the left panel, and average neural decoding accuracies are displayed on the right panel. Each subfigure has x-axis ranges from 0 to 3 seconds relative to the movement onset (*t* = 0), with 0.5 grid interval. On the left panel, the average ERPs over all trials corresponding to the same target are displayed along with their standard deviation (shaded areas). The 4 targets of Tgt1, Tgt2, Tgt3, and Tgt4 are color coded in red, green, blue, and magenta, respectively. The y-axis shows an amplitude range of 30 ± *μ*V. The right panels represent average neural decoding accuracy over the 4 targets, when 0.5-second time window was used. The x-axis segment of each horizontal line indicates the time window used for the EEG extraction for LSVM implementation, whereas the y-axis value shows the average success rates. The y-axis ranges from 0.5 to 1 of accuracy, and each vertical line represents the standard deviation over 5 folds.

The frequency range of 0.1-30 Hz was specifically applied to display neural activities correlated with hand and arm movements [39, 80–82]. In particular, movement-related cortical potential (MRCP) is known to be a unique slow EEG dynamic, containing 0.1-4 Hz components, which can be observed before and during movement in the primary motor cortex and supplementary motor cortex [39, 81, 83, 84]. In addition, event-related de/synchronization (ERD/ERS) represent movement-induced changes in EEG power within the alpha (8-13 Hz) and beta (14-30 Hz) bands, with ERD reflecting the reduction in rhythmic power during active movement phases and ERS reflecting increased power during relaxation or post-movement phases [85, 86].

Left panels of Figures 9, 10, and 11 illustrate the average ERPs on the C3 channel to visualize the characteristic neural dynamics associated with the right hand movements [87]. All figures show distinctive ERPs with respect to the 4 different targets. In general, immediately after movement onset at time 0, an increase in amplitude occurred for ERPs corresponding to Tgt3 and Tgt4 (blue and magenta lines in the left panels of Figure 9, 10, and 11) and a decrease in amplitude for Tgt1 and Tgt2 (red and green lines in the left panels of Figure 9, 10, and 11). These patterns likely reflect differences in neural processing related to the spatial location of the target objects. Although the overall patterns are similar, the scale of the amplitude varies significantly across all subjects, highlighting the well-known inter subject variability in ERPs of movement intentions [88].

### Movement Intention Classification

Our data is further validated by performing movement intention classification using a linear support vector machine (LSVM). LSVM is one of the most widely applied classifiers in EEG-based BMIs due to its simple yet effective performances [89, 90]. In our experimental paradigm, each target corresponds to a unique class, making it a 4-class classification task. This multi-class classification was performed using the one-vs-all approach [91]. To evaluate its performance, the accuracy was calculated as the proportion of the number of correctly classified targets over the total number of valid trials within a data file [92].

We used the ERP described in section Event-Related Potential (ERP) as input. The bandpass filtered EEG was segmented in 0.5 seconds, and then the segmented EEG from each channel was concatenated into a single vector for each trial, creating a vector with 3878 elements (125 samples × 31 channels). Non-overlapping sliding windows from 0 to 3 seconds was applied, resulting in 6 different time segments: 0-0.5, 0.5-1, 1-1.5, 1.5-2, 2-2.5, and 2.5-3 seconds. LSVM was trained and evaluated for each time segment, and average accuracies over 5-fold stratified cross-validation are displayed in the right panels of Figure 9, 10, and 11. These figures display the average accuracy for each recording individually.

Furthermore, when the mean accuracy of each recording with respect to each extracted window was averaged over all 49 recordings, all segmented window cases show over 75% of average accuracy (Figure 12). The peak average accuracy was observed when using the 0.5-1 second window, reaching 89.5 ± 7.6% accuracy. In addition, within the first 1.5 seconds, the average classification accuracy remains predominantly above 80% (82.8 ± 8.9% and 86.0 ± 8.4% for 0-0.5 and 1-1.5 second windows, respectively) supporting the reliability of the dataset to distinguish different movement intentions.

**Figure 12.**
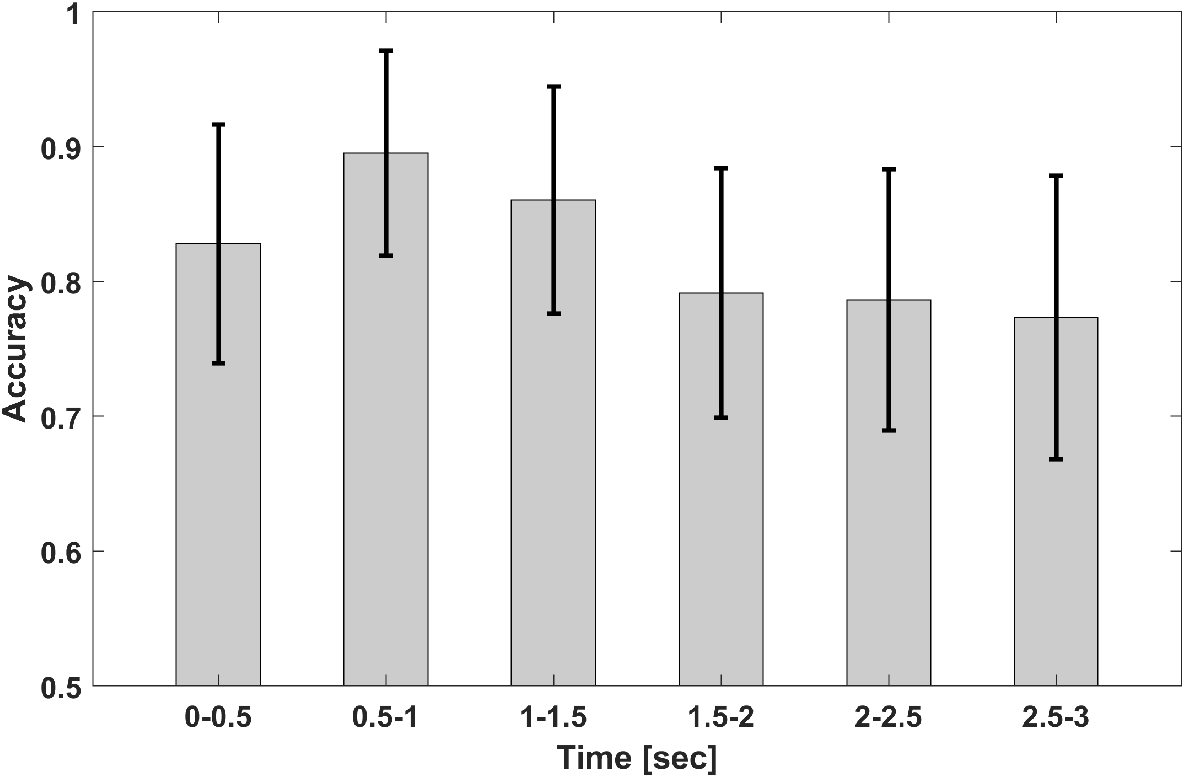
Average Accuracy Across All 49 Recordings. The bar graph represents the average neural decoding accuracy across all 49 recordings. The x-axis indicates the time window used for the EEG feature extraction, whereas the y-axis corresponds to the average accuracy. The y-axis ranges from 0.5 to 1, and each black vertical line on the bars represents the standard deviation across all 49 recordings.

## Code availability

All codes to generate the audio cues for the experiment, to process data for trial start and end and movement onset time detections, and to produce all reported results, including ERP visualization and implementation of the LSVM classifier, are accessible under *src* directory. Detailed explanations of each MATLAB code are provided in section Main Directory 2: src.

## Acknowledgments

This work was partially supported by the College of Engineering Research Mini Grant Program and Dr. Jihye Bae’s Start Up Funds provided by the Department of Electrical and Computer Engineering at the University of Kentucky.

## Author contributions statement

J. Bae designed, organized, and supervised the overall study and contributed to the manuscript writing. B. R. T. contributed to the collection and analysis, and to writing the manuscript. All provided codes are created by B. R. T. and reviewed by J. Bae. J. Boggess assisted in data collection.

## References

[1] Iahn Cajigas and Aditya Vedantam. Brain-computer interface, neuromodulation, and neurorehabilitation strategies for spinal cord injury. Neurosurgery Clinics, 32(3): 407–417, 2021. ISSN 1042-3680. doi:10.1016/j.nec.2021.03.012.

[2] Stefano Silvoni, Ander Ramos-Murguialday, Marianna Cavinato, Chiara Volpato, Giulia Cisotto, Andrea Turolla, Francesco Piccione, and Niels Birbaumer. Brain-computer interface in stroke: a review of progress. Clinical EEG and neuroscience, 42(4): 245–252, 2011. ISSN 1550-0594. doi:10.1177/155005941104200410.

[3] Theresa M Vaughan. Brain-computer interfaces for people with amyotrophic lateral sclerosis. Handbook of Clinical Neurology, 168: 33–38, 2020. ISSN 0072-9752. doi:10.1016/b978-0-444-63934-9.00004-4.

[4] Stephan Waldert. Invasive vs. non-invasive neuronal signals for brain-machine interfaces: Will one prevail? Frontiers in Neuroscience, 10, 2016. ISSN 1662-453X. doi:10.3389/fnins.2016.00295.

[5] Gerwin Schalk and Eric C Leuthardt. Brain-computer interfaces using electrocorticographic signals. IEEE reviews in biomedical engineering, 4: 140–154, 2011. ISSN 1937-3333. doi:10.1109/RBME.2011.2172408.

[6] Mark L Homer, Arto V Nurmikko, John P Donoghue, and Leigh R Hochberg. Sensors and decoding for intracortical brain computer interfaces. Annual review of biomedical engineering, 15(1): 383–405, 2013. ISSN 1523-9829. doi:10.1146/annurev-bioeng-071910-124640.

[7] Noman Naseer and Keum-Shik Hong. fnirs-based brain-computer interfaces: a review. Frontiers in human neuroscience, 9:3, 2015. ISSN 1662-5161. doi:10.3389/fnhum.2015.00172.

[8] Bettina Sorger and Rainer Goebel. Real-time fmri for brain-computer interfacing. Handbook of clinical neurology, 168: 289–302, 2020. ISSN 0072-9752.

[9] Michael L Martini, Eric Karl Oermann, Nicholas L Opie, Fedor Panov, Thomas Oxley, and Kurt Yaeger. Sensor modalities for brain-computer interface technology: a comprehensive literature review. Neurosurgery, 86(2): E108–E117, 2020. ISSN 0148-396X.

[10] Byoung-Kyong Min, Matthew J Marzelli, and Seung-Schik Yoo. Neuroimaging-based approaches in the brain–computer interface. Trends in biotechnology, 28(11): 552–560, 2010. ISSN 0167-7799.

[11] Roberto Portillo-Lara, Bogachan Tahirbegi, Christopher AR Chapman, Josef A Goding, and Rylie A Green. Mind the gap: State-of-the-art technologies and applications for eeg-based brain–computer interfaces. APL bioengineering, 5(3), 2021. doi:10.1063/5.0047237.

[12] M. Rashid, N. Sulaiman, P. P. Abdul Majeed A, R. M. Musa, A. F. Ab Nasir, B. S. Bari, and S. Khatun. Current status, challenges, and possible solutions of eeg-based brain-computer interface: A comprehensive review. Front Neurorobot, 14:25, 2020. ISSN 1662-5218 (Print) 1662-5218 (Electronic) 1662-5218 (Linking). doi:10.3389/fnbot.2020.00025.

[13] Ioulietta Lazarou, Spiros Nikolopoulos, Panagiotis C Petrantonakis, Ioannis Kompatsiaris, and Magda Tsolaki. Eeg-based brain–computer interfaces for communication and rehabilitation of people with motor impairment: a novel approach of the 21 st century. Frontiers in human neuroscience, 12:14, 2018. ISSN 1662-5161. doi:10.3389/fnhum.2018.00014.

[14] Lei Cao, Bin Xia, Oladazimi Maysam, Jie Li, Hong Xie, and Niels Birbaumer. A synchronous motor imagery based neural physiological paradigm for brain computer interface speller. Frontiers in Human Neuroscience, 11, May 2017. doi:10.3389/fnhum.2017.00274.

[15] Jiahui Pan, XueNing Chen, Nianming Ban, JiaShao He, Jiayi Chen, and Haiyun Huang. Advances in p300 brain–computer interface spellers: toward paradigm design and performance evaluation. Frontiers in Human Neuroscience, 16:1077717, 2022. ISSN 1662-5161. doi:10.3389/fnhum.2022.1077717.

[16] None Yuanqing Li, None Jiahui Pan, None Fei Wang, and None Zhuliang Yu. A hybrid bci system combining p300 and ssvep and its application to wheelchair control. IEEE Transactions on Biomedical Engineering, 60(11): 3156–3166, Oct 2013. doi:10.1109/tbme.2013.2270283.

[17] Rinku Roy, M. Mahadevappa, and C.S Kumar. Trajectory path planning of eeg controlled robotic arm using ga. Procedia Computer Science, 84:147–151, Jan 2016. doi:10.1016/j.procs.2016.04.080.

[18] M. Serdar Bascil, A. Y. Tesneli, and F. Temurtas. Multi-channel eeg signal feature extraction and pattern recognition on horizontal mental imagination task of 1-d cursor movement for brain computer interface. Australas Phys Eng Sci Med, 38(2): 229–39, 2015. ISSN 0158-9938. doi:10.1007/s13246-015-0345-6.

[19] Jinyi Long, Yuanqing Li, Tianyou Yu, and Zhenghui Gu. Target selection with hybrid feature for bci-based 2-d cursor control. IEEE Transactions on Biomedical Engineering, 59(1):132–140, Jan 2012. doi:10.1109/tbme.2011.2167718.

[20] M. Nakanishi, Y. Wang, X. Chen, Y. T. Wang, X. Gao, and T. P. Jung. Enhancing detection of ssveps for a high-speed brain speller using task-related component analysis. IEEE Trans Biomed Eng, 65(1): 104–112, 2018. ISSN 0018-9294 (Print) 0018-9294. doi:10.1109/tbme.2017.2694818.

[21] Lei Cao, Jie Li, Hongfei Ji, and Changjun Jiang. A hybrid brain computer interface system based on the neurophysiological protocol and brain-actuated switch for wheelchair control. Journal of neuroscience methods, 229: 33–43, 2014. ISSN 0165-0270.

[22] Rihab Bousseta, I El Ouakouak, M Gharbi, and F Regragui. Eeg based brain computer interface for controlling a robot arm movement through thought. Irbm, 39(2): 129–135, 2018. ISSN 1959-0318.

[23] M. K. Lu, N. Arai, C. H. Tsai, and U. Ziemann. Movement related cortical potentials of cued versus self-initiated movements: double dissociated modulation by dorsal premotor cortex versus supplementary motor area rtms. Hum Brain Mapp, 33(4): 824–39, 2012. ISSN 1065-9471 (Print) 1065-9471. doi:10.1002/hbm.21248.

[24] A. M. Savic, I. K. Niazi, and M. B. Popovic. Self-paced vs. cue-based motor task: The difference in corti-cal activity. In 2011 19thTelecommunications Forum (TELFOR) Proceedings of Papers, pages 39–42, 2011. doi:10.1109/TELFOR.2011.6143887.

[25] S Jankelowitz and J Colebatch. Movement-related potentials associated with self-paced, cued and imagined arm movements. Experimental brain research, 147: 98–107, 2002. ISSN 0014-4819.

[26] Murat Kaya, Mustafa Kemal Binli, Erkan Ozbay, Hilmi Yanar, and Yuriy Mishchenko. A large electroencephalographic motor imagery dataset for electroencephalographic brain computer interfaces. Scientific Data, 5(1): 180211, 2018. ISSN 2052-4463. doi:10.1038/sdata.2018.211.

[27] Benjamin Blankertz, Guido Dornhege, Matthias Krauledat, Klaus-Robert Müller, and Gabriel Curio. The non-invasive berlin brain–computer interface: Fast acquisition of effective performance in untrained subjects. NeuroImage, 37(2), 2007. ISSN 1053-8119. doi:10.1016/j.neuroimage.2007.01.051.

[28] Clemens Brunner, Robert Leeb, Gernot Müller-Putz, Alois Schlögl, and Gert Pfurtscheller. Bci competition 2008–graz data set a. Institute for knowledge discovery (laboratory of brain-computer interfaces), Graz University of Technology, 16: 1–6, 2008.

[29] Robert Leeb, Clemens Brunner, G Müller-Putz, A Schlögl, and GJGUOT Pfurtscheller. Bci competition 2008–graz data set b. Graz University of Technology, Austria, 16: 1–6, 2008.

[30] Jianjun Meng, Shuying Zhang, Angeliki Bekyo, Jaron Olsoe, Bryan Baxter, and Bin He. Noninvasive electroencephalogram based control of a robotic arm for reach and grasp tasks. Scientific Reports, 6(1): 38565, 2016. ISSN 2045-2322. doi:10.1038/srep38565.

[31] Patrick Ofner, Andreas Schwarz, Joana Pereira, and Gernot R. Müller-Putz. Upper limb movements can be decoded from the time-domain of low-frequency eeg. PLOS ONE, 12(8):e0182578, 2017. doi:10.1371/journal.pone.0182578.

[32] Hohyun Cho, Minkyu Ahn, Sangtae Ahn, Kwon Moonyoung, and Sung Chan Jun. Supporting data for “eeg datasets for motor imagery brain computer interface”, 2017.

[33] Xuelin Ma, Shuang Qiu, and Huiguang He. Multi-channel eeg recording during motor imagery of different joints from the same limb. Scientific Data, 7(1): 191, 2020. ISSN 2052-4463. doi:10.1038/s41597-020-0535-2.

[34] Ji-Hoon Jeong, Jeong-Hyun Cho, Kyung-Hwan Shim, Byoung-Hee Kwon, Byeong-Hoo Lee, Do-Yeun Lee, Dae-Hyeok Lee, and Seong-Whan Lee. Multimodal signal dataset for 11 intuitive movement tasks from single upper extremity during multiple recording sessions. GigaScience, 9(10), 2020. ISSN 2047-217X. doi:10.1093/gigascience/giaa098.

[35] S. Rasheed and W. Mumtaz. Classification of hand-grasp movements of stroke patients using eeg data. In 2021 International Conference on Artificial Intelligence (ICAI), pages 86–90. doi:10.1109/ICAI52203.2021.9445231.

[36] Haijie Liu, Penghu Wei, Haochong Wang, Xiaodong Lv, Wei Duan, Meijie Li, Yan Zhao, Qingmei Wang, Xinyuan Chen, Gaige Shi, Bo Han, and Junwei Hao. An eeg motor imagery dataset for brain computer interface in acute stroke patients. Scientific Data, 11(1): 131, 2024. ISSN 2052-4463. doi:10.1038/s41597-023-02787-8.

[37] Gert Pfurtscheller, Clemens Brunner, Alois Schlögl, and FH Lopes Da Silva. Mu rhythm (de) synchronization and eeg single-trial classification of different motor imagery tasks. NeuroImage, 31(1): 153–159, 2006. ISSN 1053-8119.

[38] Weibo Yi, Shuang Qiu, Hongzhi Qi, Lixin Zhang, Baikun Wan, and Dong Ming. Eeg feature comparison and classification of simple and compound limb motor imagery. Journal of neuroengineering and rehabilitation, 10: 1–12, 2013.

[39] J. Wang, L. Bi, and W. Fei. Eeg-based motor bcis for upper limb movement: Current techniques and future insights. IEEE Trans Neural Syst Rehabil Eng, 31: 4413–4427, 2023. ISSN 1558-0210 (Electronic) 1534-4320 (Linking). doi:10.1109/TNSRE.2023.3330500.

[40] M Ebrahim M Mashat, Chin-Teng Lin, and Dingguo Zhang. Effects of task complexity on motor imagery-based brain–computer interface. IEEE Transactions on Neural Systems and Rehabilitation Engineering, 27(10): 2178–2185, 2019. ISSN 1534-4320.

[41] Lili Li, Jing Wang, Guanghua Xu, Min Li, and Jun Xie. The study of object-oriented motor imagery based on eeg suppression. PLoS One, 10(12):e0144256, 2015. ISSN 1932-6203.

[42] Attila Korik, Ronen Sosnik, Nazmul Siddique, and Damien Coyle. Decoding imagined 3d hand movement trajectories from eeg: evidence to support the use of mu, beta, and low gamma oscillations. Frontiers in neuroscience, 12:130, 2018. ISSN 1662-453X.

[43] Han Yuan, Christopher Perdoni, and Bin He. Relationship between speed and eeg activity during imagined and executed hand movements. Journal of neural engineering, 7(2): 026001, 2010. ISSN 1741-2552.

[44] Austin J. Hurst and Shaun G. Boe. Imagining the way forward: A review of contemporary motor imagery theory. Frontiers in Human Neuroscience, 16, 2022. ISSN 1662-5161. doi:10.3389/fnhum.2022.1033493.

[45] Yvonne Höller, Jürgen Bergmann, Martin Kronbichler, Julia Sophia Crone, Elisabeth Verena Schmid, Aljoscha Thomschewski, Kevin Butz, Verena Schütze, Peter Höller, and Eugen Trinka. Real movement vs. motor imagery in healthy subjects. International Journal of Psychophysiology, 87(1): 35–41, 2013. ISSN 0167-8760.

[46] M. T. Carrillo-de-la Peña, S. Galdo-Álvarez, and C. Lastra-Barreira. Equivalent is not equal: Primary motor cortex (mi) activation during motor imagery and execution of sequential movements. Brain Research, 1226: 134–143, 2008. ISSN 0006-8993. doi:10.1016/j.brainres.2008.05.089.

[47] Helen O’Shea and Aidan Moran. Does motor simulation theory explain the cognitive mechanisms underlying motor imagery? a critical review. Frontiers in Human Neuroscience, 11, 2017. ISSN 1662-5161. doi:10.3389/fnhum.2017.00072.

[48] Matthew D. Luciw, Ewa Jarocka, and Benoni B. Edin. Multi-channel eeg recordings during 3,936 grasp and lift trials with varying weight and friction. Scientific Data, 1(1): 140047, 2014. ISSN 2052-4463. doi:10.1038/sdata.2014.47.

[49] Andreea I. Sburlea and Gernot R. Müller-Putz. Exploring representations of human grasping in neural, muscle and kinematic signals. Scientific Reports, 8(1), Nov 2018. doi:10.1038/s41598-018-35018-x.

[50] Andreas Schwarz, Carlos Escolano, Luis Montesano, and Gernot R. Müller-Putz. Analyzing and decoding natural reach-and-grasp actions using gel, water and dry eeg systems. Frontiers in Neuroscience, 14, 2020. ISSN 1662-453X. doi:10.3389/fnins.2020.00849.

[51] Fabien Lotte, Camille Jeunet, Ricardo Chavarriaga, Laurent Bougrain, Dave E Thompson, Reinhold Scherer, Md Rakibul Mowla, Andrea Kübler, Moritz Grosse-Wentrup, and Karen Dijkstra. Turning negative into positives! exploiting ‘negative’results in brain–machine interface (bmi) research. Brain-Computer Interfaces, 6(4): 178–189, 2019. ISSN 2326-263X. doi:10.1080/2326263X.2019.1697143.

[52] F. Lotte, L. Bougrain, A. Cichocki, M. Clerc, M. Congedo, A. Rakotomamonjy, and F. Yger. A review of classification algorithms for eeg-based brain-computer interfaces: a 10 year update. J Neural Eng, 15(3): 031005, 2018. doi:10.1088/1741-2552/aab2f2.

[53] Nikunj A. Bhagat, Anusha Venkatakrishnan, Berdakh Abibullaev, Edward J. Artz, Nuray Yozbatiran, Amy A. Blank, James French, Christof Karmonik, Robert G. Grossman, Marcia K. O’Malley, Gerard E. Francisco, and Jose L. Contreras-Vidal. Design and optimization of an eeg-based brain machine interface (bmi) to an upper-limb exoskeleton for stroke survivors. Frontiers in Neuroscience, 10, 2016. ISSN 1662-453X. doi:10.3389/fnins.2016.00122.

[54] T. P. Jung, S. Makeig, C. Humphries, T. W. Lee, M. J. McKeown, V. Iragui, and T. J. Sejnowski. Removing electroencephalographic artifacts by blind source separation. Psychophysiology, 37(2): 163–78, 2000. ISSN 0048-5772 (Print) 0048-5772. doi:10.1111/1469-8986.3720163.

[55] Gabriele Gratton, Michael G.H Coles, and Emanuel Donchin. A new method for off-line removal of ocular artifact. Electroencephalography and Clinical Neurophysiology, 55(4):468–484, Apr 1983. doi:10.1016/0013-4694(83)90135-9.

[56] Xiao Jiang, Gui-Bin Bian, and Zean Tian. Removal of artifacts from eeg signals: A review. Sensors (Basel, Switzerland), 19(5): 987, 2019. ISSN 1424-8220. doi:10.3390/s19050987.

[57] Shailaja Kotte and J. R. K. Kumar Dabbakuti. Methods for removal of artifacts from eeg signal: A review. Journal of Physics: Conference Series, 1706(1): 012093, 2020. ISSN 1742-6596 1742-6588. doi:10.1088/1742-6596/1706/1/012093.

[58] Rakesh Ranjan, Bikash Chandra Sahana, and Ashish Kumar Bhandari. Ocular artifact elimination from electroencephalography signals: A systematic review. Biocybernetics and Biomedical Engineering, 41(3): 960–996, 2021. ISSN 02085216. doi:10.1016/j.bbe.2021.06.007.

[59] Rodney J Croft, Jody S Chandler, Robert J Barry, Nicholas R Cooper, and Adam R Clarke. Eog correction: a comparison of four methods. Psychophysiology, 42(1): 16–24, 2005. ISSN 0048-5772. doi:10.1111/j.1468-8986.2005.00264.x.

[60] Kevin Hooks, Refaat El-Said, and Qiushi Fu. Decoding reach-to-grasp from eeg using classifiers trained with data from the contralateral limb. Frontiers in Human Neuroscience, 17:1302647, 2023. ISSN 1662-5161.

[61] Mehmet Dursun, Seral Özsen, Cüneyt Yücelbas, Sule Yücelbas, Gülay Tezel, Serkan Küççüktürk, and Sebnem Yosunkaya. A new approach to eliminating eog artifacts from the sleep eeg signals for the automatic sleep stage classification. Neural Computing and Applications, 28: 3095–3112, 2017. ISSN 0941-0643.

[62] Jin Wu, Jiacai Zhang, and Li Yao. An automated detection and correction method of eog artifacts in eeg-based bci. In 2009 ICME International Conference on Complex Medical Engineering, pages 1–5. IEEE. ISBN 1424433150.

[63] Lüder Deecke. Planning, preparation, execution, and imagery of volitional action. Cognitive Brain Research, 3 (2):59–64, 1996. ISSN 0926-6410. doi:10.1016/0926-6410(95)00046-1.

[64] L. M. Rueda-Delgado, E. Solesio-Jofre, D. J. Serrien, D. Mantini, A. Daffertshofer, and S. P. Swinnen. Understanding bimanual coordination across small time scales from an electrophysiological perspective. Neuroscience Biobehavioral Reviews, 47: 614–635, 2014. ISSN 0149-7634. doi:10.1016/j.neubiorev.2014.10.003.

[65] G. R. Muller-Putz, R. J. Kobler, J. Pereira, C. Lopes-Dias, L. Hehenberger, V. Mondini, V. Martinez-Cagigal, N. Srisrisawang, H. Pulferer, L. Batistic, and A. I. Sburlea. Feel your reach: An eeg-based framework to continuously detect goal-directed movements and error processing to gate kinesthetic feedback informed artificial arm control. Front Hum Neurosci, 16:841312, 2022. ISSN 1662-5161 (Print) 1662-5161(Electronic) 1662-5161 (Linking). doi:10.3389/fnhum.2022.841312.

[66] Chang-Hee Han, Klaus-Robert Müller, and Han-Jeong Hwang. Brain-switches for asynchronous brain–computer interfaces: A systematic review. Electronics, 9(3), 2020. ISSN 2079-9292. doi:10.3390/electronics9030422.

[67] G. Pfurtscheller, C. Neuper, G.R Muller, B Obermaier, G Krausz, A. Schlogl, R Scherer, B. Graimann, C. Keinrath, D. Skliris, M Wortz, G Supp, and C Schrank. Graz-bci: state of the art and clinical applications. IEEE Transactions on Neural Systems and Rehabilitation Engineering, 11(2):1–4, Jun 2003.

[68] Sura Rodpongpun, Thapanan Janyalikit, and Chotirat Ann Ratanamahatana. Influential factors of an asynchronous bci for movement intention detection. Computational and Mathematical Methods in Medicine, 2020:8573754, 2020. ISSN 1748-670X. doi:10.1155/2020/8573754.

[69] Shiqi Sun, Xiya Cao, and Qining Wang. Continuous decoding of movement onset and offset of sustained movements from cortical activities. In 2016 IEEE/SICE International Symposium on System Integration (SII), pages 809–814. IEEE, 2016. doi:10.1109/SII.2016.7844099.

[70] Bhoj Raj Thapa, John Boggess, and Jihye Bae. A large electroencephalogram database of freewill reaching and grasping tasks for brain machine interfaces. figshare, Mar 2025. doi:10.6084/m9.figshare.28632599.

[71] Pauli Virtanen, Ralf Gommers, Travis E. Oliphant, Matt Haberland, Tyler Reddy, David Cournapeau, Evgeni Burovski, Pearu Peterson, Warren Weckesser, Jonathan Bright, Stéfan J. van der Walt, Matthew Brett, Joshua Wilson, K. Jarrod Millman, Nikolay Mayorov, Andrew R. J. Nelson, Eric Jones, Robert Kern, Eric Larson, C J Carey, Ilhan Polat, Yu Feng, Eric W. Moore, Jake VanderPlas, Denis Laxalde, Josef Perktold, Robert Cimrman, Ian Henriksen, E. A. Quintero, Charles R. Harris, Anne M. Archibald, Antônio H. Ribeiro, Fabian Pedregosa, Paul van Mulbregt, and SciPy 1.0 Contributors. SciPy 1.0: Fundamental Algorithms for Scientific Computing in Python. Nature Methods, 17: 261–272, 2020. doi:10.1038/s41592-019-0686-2.

[72] Manousos A. Klados, Christos Papadelis, Christoph Braun, and Panagiotis D. Bamidis. Reg-ica: A hybrid methodology combining blind source separation and regression techniques for the rejection of ocular artifacts. Biomedical Signal Processing and Control, 6(3): 291–300, 2011. ISSN 1746-8094. doi:10.1016/j.bspc.2011.02.001.

[73] Erkki Oja and Zhijian Yuan. The fastica algorithm revisited: Convergence analysis. IEEE transactions on Neural Networks, 17(6): 1370–1381, 2006. ISSN 1045-9227.

[74] T. W. Lee, M. Girolami, and T. J. Sejnowski. Independent component analysis using an extended infomax algorithm for mixed subgaussian and supergaussian sources. Neural Computation, 11(2): 417–441, 1999. ISSN 0899-7667. doi:10.1162/089976699300016719.

[75] Y. Tang, J. Tang, and A. Gong. Removal of ocular artifact from eeg using jade. In 2007 1st International Conference on Bioinformatics and Biomedical Engineering, pages 566–569. ISBN 2151-7622. doi:10.1109/ICBBE.2007.148.

[76] Jose Antonio Urigüen and Begoña Garcia-Zapirain. Eeg artifact removal—state-of-the-art and guidelines. Journal of neural engineering, 12(3): 031001, 2015. ISSN 1741-2552. doi:10.1088/1741-2560/12/3/031001.

[77] S. Romero, M. A. Mananas, and M. J. Barbanoj. A comparative study of automatic techniques for ocular artifact reduction in spontaneous eeg signals based on clinical target variables: a simulation case. Comput Biol Med, 38(3): 348–60, 2008. ISSN 0010-4825 (Print) 0010-4825 (Linking). doi:10.1016/j.compbiomed.2007.12.001.

[78] Laurent Albera, Amar Kachenoura, Pierre Comon, Ahmad Karfoul, Fabrice Wendling, Lotfi Senhadji, and Isabelle Merlet. Ica-based eeg denoising: a comparative analysis of fifteen methods. Bulletin of the Polish Academy of Sciences: Technical Sciences, 60(3 Special issue on Data Mining in Bioengineering):407–418, 2012.

[79] Markus Plank. Ocular correction ica, 2022. https://www.brainproducts.com/support-resources/ocular-correction-ica/.

[80] Simona Noviello, Saman Kamari Songhorabadi, Zhiqing Deng, Chao Zheng, Juan Chen, Angelo Pisani, Elena Franchin, Enrica Pierotti, Elena Tonolli, Simona Monaco, Louis Renoult, and Irene Sperandio. Temporal features of size constancy for perception and action in a real-world setting: A combined eeg-kinematics study. Neuropsychologia, 193:108746, 2024. ISSN 0028-3932. doi:10.1016/j.neuropsychologia.2023.108746.

[81] Eileen Lew, Ricardo Chavarriaga, Stefano Silvoni, and José del R. Millán. Detection of self-paced reaching movement intention from eeg signals. Frontiers in Neuroengineering, 5(13), 2012. ISSN 1662-6443. doi:10.3389/fneng.2012.00013.

[82] C. C. Kuo, W. S. Lin, C. A. Dressel, and A. W. L. Chiu. Classification of intended motor movement using surface eeg ensemble empirical mode decomposition. In 2011 Annual International Conference of the IEEE Engineering in Medicine and Biology Society, pages 6281–6284. ISBN 1558-4615. doi:10.1109/IEMBS.2011.6091550.

[83] Aqsa Shakeel, Muhammad Samran Navid, Muhammad Nabeel Anwar, Suleman Mazhar, Mads Jochumsen, and Imran Khan Niazi. A review of techniques for detection of movement intention using movement-related cortical potentials. Computational and Mathematical Methods in Medicine, 2015:346217, 2015. ISSN 1748-670X. doi:10.1155/2015/346217.

[84] James G Colebatch. Bereitschaftspotential and movement-related potentials: origin, significance, and application in disorders of human movement. Movement Disorders, 22(5): 601–610, 2007. ISSN 0885-3185.

[85] Hao Gu, Jian Wang, Fengyuan Jiao, Yan Han, Wang Xu, and Xin Zhao. Decoding electroencephalography underlying natural grasp tasks across multiple dimensions. Electronics, 12(18), 2023. ISSN 2079-9292. doi:10.3390/electronics12183894.

[86] Gert Pfurtscheller and FH Lopes Da Silva. Event-related eeg/meg synchronization and desynchronization: basic principles. Clinical neurophysiology, 110(11): 1842–1857, 1999. ISSN 1388-2457.

[87] A. M. Savic, E. R. Lontis, N. Mrachacz-Kersting, and M. B. Popovic. Dynamics of movement-related cortical potentials and sensorimotor oscillations during palmar grasp movements. Eur J Neurosci, 51(9): 1962–1970, 2020. ISSN 1460-9568 (Electronic) 0953-816X (Linking). doi:10.1111/ejn.14629.

[88] Marta Borràs, Sergio Romero, Joan F. Alonso, Alejandro Bachiller, Leidy Y. Serna, Carolina Migliorelli, and Miguel A. Mañanas. Influence of the number of trials on evoked motor cortical activity in eeg recordings. Journal of Neural Engineering, 19(4): 046050, 2022. doi:10.1088/1741-2552/ac86f5.

[89] Saim Rasheed. A review of the role of machine learning techniques towards brain–computer interface applications. Machine Learning and Knowledge Extraction, 3(4): 835–862, 2021. ISSN 2504-4990. doi:10.3390/make3040042.

[90] Maham Saeidi, Waldemar Karwowski, Farzad V. Farahani, Krzysztof Fiok, Redha Taiar, P. A. Hancock, and Awad Al-Juaid. Neural decoding of eeg signals with machine learning: A systematic review. Brain Sciences, 11(11): 1525, 2021. ISSN 2076-3425.

[91] Ryan Rifkin and Aldebaro Klautau. In defense of one-vs-all classification. The Journal of Machine Learning Research, 5: 101–141, 2004. ISSN 1532-4435.

[92] Mohammad Hossin and Md Nasir Sulaiman. A review on evaluation metrics for data classification evaluations. International journal of data mining knowledge management process, 5(2): 1, 2015. ISSN 2231-007X. doi:10.5121/ijdkp.2015.5201.

