## Supplementary Table 1 for "A Large Electroencephalogram Database of Freewill Reaching and Grasping Tasks for Brain Machine Interfaces"

### List of Tables

Supplementary Table 1. List of publicly available EEG datasets for upper limb movement……………..3

The table (Supplementary Table 1) details critical information on the datasets, including authors, publication years, participant demographics, the number of EEG channels, where freewill or actual movement execution was involved, continuous EEG data availability, EOG availability, and experimental paradigms. The last two rows are colored with a grey background provide datasets that perform freewill tasks. The citations here correspond to the reference in the data descriptor.

Supplementary Table 1. List of publicly available EEG datasets for upper limb movement

| Author | Year | # of Subjects | Age [years] | Health Conditions | # of EEG Channels | Freewill | Imagery/ Execution | Continuous EEG | EOG | Experimental Paradigm | | |
|---|---|---|---|---|---|---|---|---|---|---|---|---|
| | | | | | | | | | | Task | # of Classes | Task Details |
| B. Blankertz, et al.[27] | 2007 | 10 | 26~46 | Healthy | 59 | No | Imagery a) | **Yes** | **Yes** | Hand and leg imagery | 3 | Left-hand, right-hand, and right-foot imagery guided by visual cues |
| C. Brunner, et al.[28] | 2008 | 9 | N/A | Healthy | 22 | No | Imagery a) | **Yes** | **Yes** | Hand, feet, and tongue imagery | 4 | Left-hand, right-hand, both feet, and tongue movement imagery guided by visual cues |
| R. Leeb, et al.[29] | 2008 | 9 | N/A | Healthy | 3 | No | Imagery a) | **Yes** | **Yes** | Left- and right-hand motor imagery | 2 | Left- and right-hand motor imagery |
| J. Meng, et al.[30] | 2016 | 13 | 18~54 | Healthy | 64 | No | Imagery a) | **Yes** | No | Left and right-hand motor imagery | 4 | Left hand, right hand, both hands, and relaxation imagery for left, right, up and down movements |
| P. Ofner, et al.[31] | 2017 | 15 | 27±5 | Healthy | 61 | No | Imagery b) | **Yes** | No | Movements of the right upper limb | 6 | Elbow flexion/extension, forearm supination/pronation, hand opening/closing |
| H. Cho, et al.[32] | 2017 | 52 | 24.8± 3.86 | Healthy | 64 | No | Imagery b) | **Yes** | **Yes** | Left-hand and right-hand movement | 4 | Touching each index, middle, ring, and little finger to the thumb with both hands. |
| M. Kaya, et al.[26] | 2018 | 8 | 20~35 | Healthy | 19 | No | Imagery b) | **Yes** | No | Finger movement imagery | 5 | Five fingers flexion imagery of a single hand |
| M. Kaya, et al.[26] | 2018 | 7 | 20~35 | Healthy | 19 | No | Imagery b) | **Yes** | No | Left- and right-hand motor imagery | 2 | Closing and opening of the fist for left-hand and right-hand motor imagery |
| M. Kaya, et al.[26] | 2018 | 12 | 20~35 | Healthy | 19 | No | Imagery a) | **Yes** | No | Hand, leg, tongue imagery | 6 | Left-hand, right-hand, left-leg, right-leg, tongue, and passive movement imagery |
| X. Ma, et al.[33] | 2020 | 25 | 19~27 | Healthy | 64 | No | Imagery a) | No | **Yes** | Right upper limb imagery | 3 | Right-hand and right-elbow imagery and resting with eyes open |
| J. Jeong, et al.[34] | 2020 | 25 | 24~32 | Healthy | 60 | No | Imagery b) | **Yes** | **Yes** | Upper extremity arm-reaching and hand-grasping task | 11 | Arm reaching in 6 directions, and hand-grasping of 3 objects (card, ball, and cup) |
| S. Rasheed, et al.[35] | 2021 | 10 | N/A | Hemiparetic stroke patients | 12 | No | Imagery b) | No | No | Left- and right-hand motor imagery | 2 | Left- and right-hand grasping imagery |
| H. Liu, et al.[36] | 2024 | 50 | 56.7 ±10.57 | Acute stroke patients | 30 | No | Imagery b) | No | **Yes** | Left- and right-hand motor imagery | 2 | Grasping a spherical object imagery with left- and right-hand |
| M. D. Luciw, et al.[48] | 2014 | 12 | 19~35 | Healthy | 32 | No | Execution | **Yes** | No | Reach and grasp-and-lift task using thumb and index finger | 6 | 6 objects in combination of 3 different weights (165, 330, or 660g) and various surface friction conditions (sandpaper, suede, or silk surface) |
| P. Ofner, et al.[31] | 2017 | 15 | 27±5 | Healthy | 61 | No | Execution | **Yes** | No | Movements of the right upper limb | 6 | Elbow flexion/extension, forearm supination/pronation, hand opening/closing |

| | | | | | | | | | | | | |
|---|---|---|---|---|---|---|---|---|---|---|---|---|
| H. Cho, et al.[32] | 2017 | 52 | 24.8 ±3.86 | Healthy | 64 | No | Execution | **Yes** | **Yes** | Left-hand and right-hand movement | 4 | Touching each index, middle, ring, and little finger to the thumb with both hands. |
| A. L. Sburlea, et al.[49] | 2018 | 31 | 25.2 ±3.4 | Healthy | 61 | No | Execution | No | **Yes** | Reach and grasp | 33 | 33 different grasp movements considering position of the thumb relative to the palm and object's shape and size |
| J. Jeong, et al.[34] | 2020 | 25 | 24~32 | Healthy | 60 | No | Execution | **Yes** | **Yes** | Upper extremity arm-reaching and hand-grasping task | 11 | Arm reaching in 6 directions, and hand-grasping of 3 objects (card, ball, and cup) |
| A. Schwarz, et al.[50] | 2020 | 45 | 15~30 | Healthy | 61 | No | Execution | **Yes** | **Yes** | Reach and grasp | 2 | Palmar grasp and lateral grasp of an empty jar, and a jar with a spoon stuck in it |
| M. Kaya, et. al.[26] | 2018 | 2 | 20~35 | Healthy | 19 | **Yes** | Execution | **Yes** | No | Key pressing | 2 | 'd' key press with left hand and 'l' key press with right hand |
| **Our Dataset** | 2025 | 23 | 20.7 ±1.8 | Healthy | 31 | **Yes** | Execution | **Yes** | **Yes** | Reach and grasp | 4 | Reach and grasp one of 4 cups with right hand and arm, where two cups are filled with water and 2 are empty |

N/A: The data is not available

Imagery a) imagining different body parts

Imagery b) imagining the actual movement execution
